# Breaking Balance: Encoding local error signals in perturbations of excitation-inhibition balance

**DOI:** 10.1101/2025.05.12.653626

**Authors:** Julian Rossbroich, Friedemann Zenke

## Abstract

Efficient learning algorithms such as backpropagation, predictive coding, and self-supervised learning require instructive learning signals to influence synaptic plasticity locally at specific neurons. This requirement is inconsistent with classic Hebbian theories and neuromodulation involving a non-specific third factor, thus raising the question of how neurobiology can achieve efficient learning. Here, we propose a simple and biologically plausible solution: local deviations from precise excitation/inhibition (E/I) balance encode error signals that instruct synaptic plasticity. Using a computational model derived from an adaptive control theory framework, we demonstrate that breaking E/I balance through targeted feedback to inhibitory interneurons, can produce neuron- or assembly-specific error signals that enable learning in multi-layer networks. Simulations reveal that such a balance-controlled plasticity mechanism is consistent with phenomenological local plasticity models while enabling online learning in hierarchically organized networks. Furthermore, we demonstrate that this framework is consistent with key features of disinhibitory microcircuit dynamics during in-vivo learning experiments. These results suggest that the brain may exploit E/I balance not only for stability, but also as a substrate for error-driven learning.

## Introduction

A central question in neuroscience is how the brain learns complex tasks across distributed networks of interacting brain areas. Machine learning algorithms that solve complex tasks require local error signals— neuron-specific cues that determine the sign of synaptic changes. Similarly, bio-plausible credit assignment algorithms, which determine which synapses to modify, and how, to improve performance on behavioral tasks, explicitly require local error signals that control the sign of plasticity [1, 2]. Moreover, neuron-specific errors, or teaching signals, are also required in self-supervised learning [3, 4] and predictive coding models [5–11]. While local error signals are fundamental to learning in hierarchical networks, the biological mechanisms for computing local errors in the brain remain elusive [2, 12, 13].

Existing models assume that neurons separate error signals from ongoing activity by utilizing segregated dendritic compartments (Fig. 1; [6–9, 11, 14–16]) or by alternating between distinct temporal phases [3, 7, 9, 14, 17–19]. However, direct experimental evidence for electrotonic segregation required for such compartmentalized schemes is limited to certain neuron types, such as cortical Layer 5 pyramidal cells [16], and, in some cases, inconsistent with phenomenological plasticity models [20–27], which raises questions about their generality.

**Figure 1.**
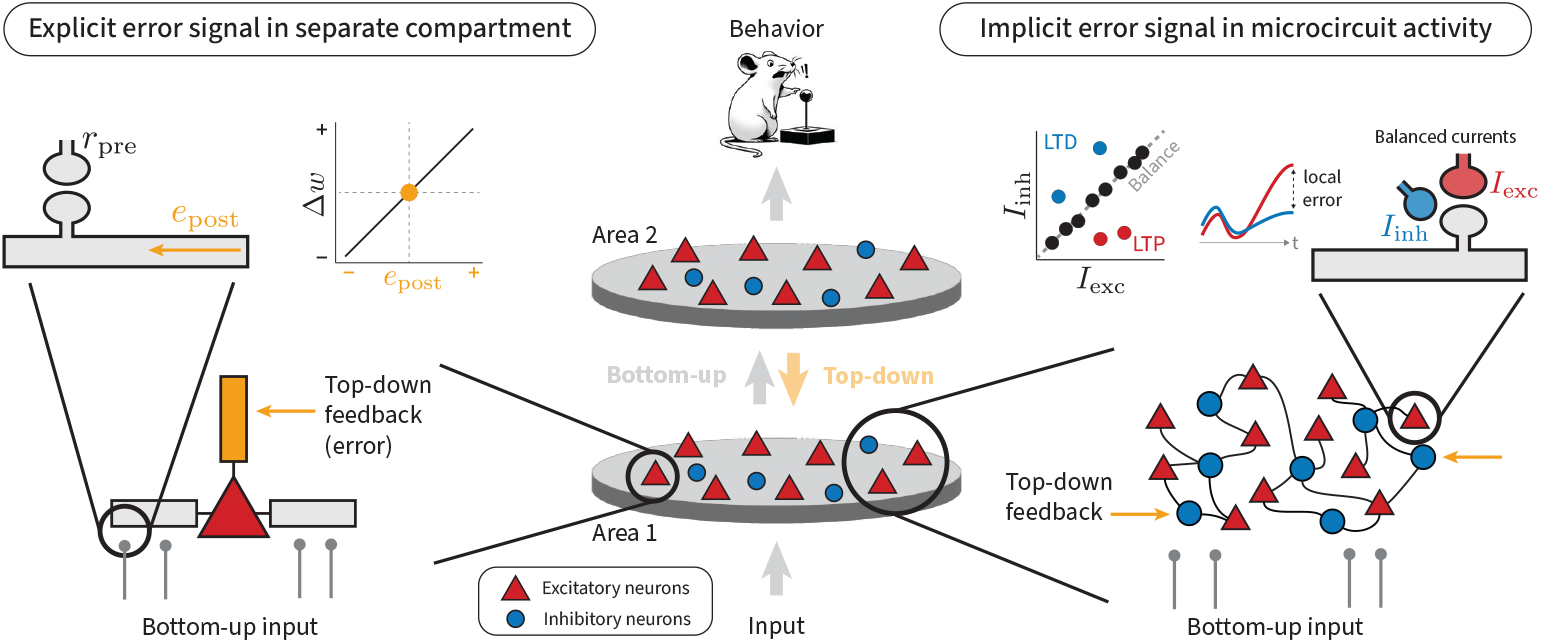
Efficient learning algorithms require local error signals that determine the sign of synaptic plasticity. How synaptic plasticity is coordinated across neuronal circuits across the brain to tune individual synapses for specific computational tasks (middle) remains a fundamental question in neuroscience. Efficient learning algorithms require local neuron-specific error signals that dictate the sign of synaptic weight changes. Most theoretically motivated models of biologically plausible credit assignment and predictive coding assume that such error signals are explicitly represented in separate temporal phases of neuronal activity or distinct neuronal compartments (left). However, such compartmentalization is often at odds with neurophysiology and inconsistent with phenomenological plasticity rules. Here we develop an alternative model that sidesteps these issues by encoding error signals in deviations from E/I balance in local E/I assemblies (right).

In contrast, E/I balance is a pervasive phenomenon widely observed in cortical circuits, whereby neurons maintain a tight balance between excitatory and inhibitory currents across stimuli and time [28–34]. While balance is considered essential to support network stability, efficient coding, and precise spike timing [35–37], its overall function remains elusive. Importantly, inhibition has been shown to influence synaptic plasticity [38–45] and learning outcomes [46–54], suggesting a potential role for E/I balance in learning beyond mere stabilization [55–57]. Nevertheless, existing work typically considered inhibition, or the lack thereof, as a permissive signal for synaptic plasticity.

This article demonstrates how E/I balance affords neuron-specific error encoding, allowing it to act as an instructive signal that could support powerful learning algorithms in the brain for solving complex tasks. Specifically, we introduce a theoretical framework in which local error signals are encoded implicitly at the microcircuit-level as controlled deviations from E/I balance (Fig. 1). We show how top-down feedback pathways with neuron-specific targeting can give rise to such systematic deviations in E/I balance and, combined with a suitable credit assignment strategy, orchestrate learning in multi-layer networks. Importantly, encoding local error signals in E/I balance constitutes a versatile circuit mechanism that reconciles powerful learning algorithms with established Hebbian principles, without the need for electronically separate dendritic compartments or distinct learning phases. By integrating error encoding with the notion of E/I balance, our model connects theories of biologically plausible credit assignment, predictive coding, and self-supervised learning with experimentally observed plasticity mechanisms, providing a fresh view into how the brain may orchestrate synaptic plasticity across large distributed neural networks to solve complex tasks.

## Results

To study whether deviations from precise E/I balance [35] can encode target-specific error signals, we consider an E/I assembly as the fundamental computational unit (Fig. 2a; Methods). We further distinguish between two types of distinct inhibitory interneuron populations (Supplementary Fig. S1). Type 1 interneurons provide feedforward inhibition and contribute to neuronal selectivity. In contrast, Type 2 interneurons are reciprocally connected and co-tuned with excitatory neurons resulting in a precise E/I balance (Fig. 2b). Signatures of such co-tuning exist in neurobiology [28, 30, 31, 33, 58] and numerous computational studies demonstrated how precise balance can emerge through suitable Hebbian plasticity mechanisms [25, 57, 59–62].

**Figure 2.**
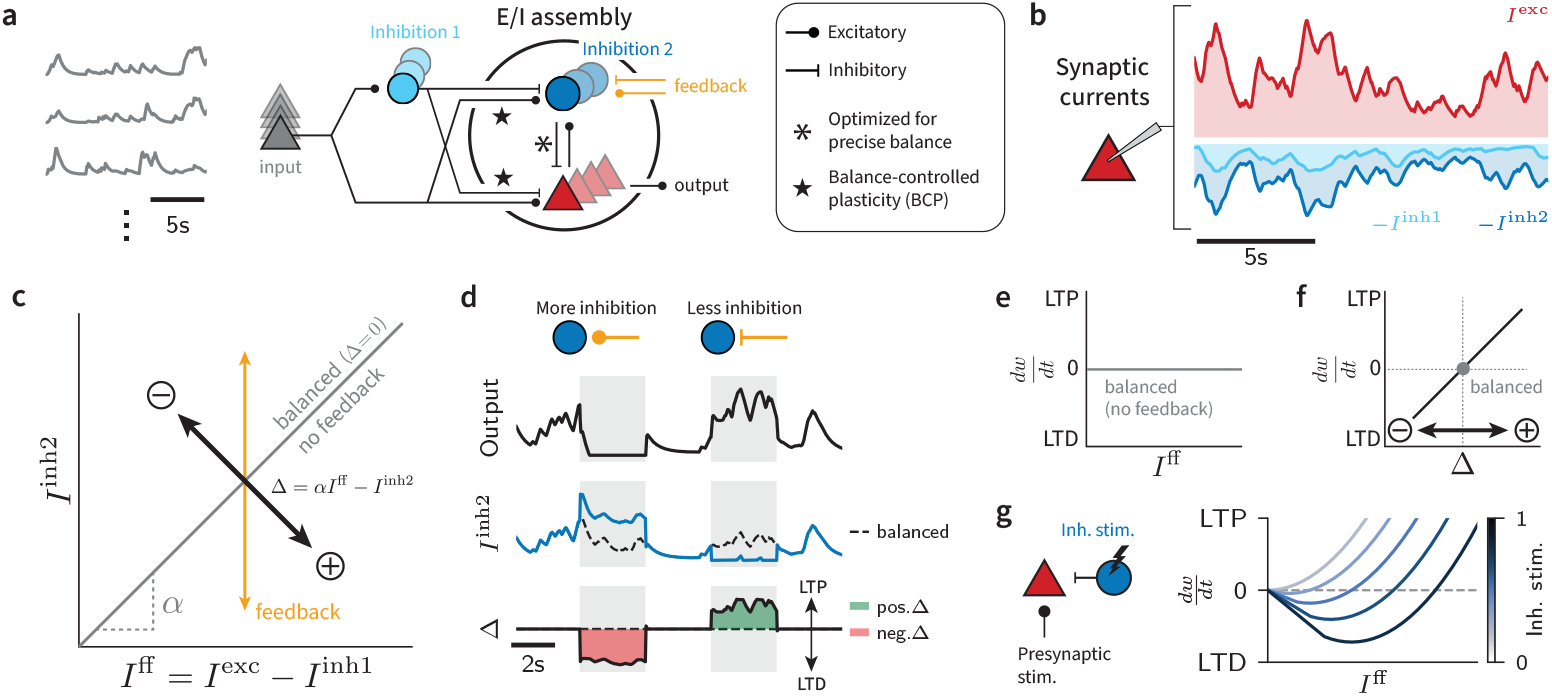
Deviations from E/I balance can encode assembly-specific error signals. **(a)** Schematic representation of an E/I assembly receiving excitatory input and feed-forward inhibition (Type 1). All input synapses are subject to balance-controlled plasticity (BCP). Within the assembly, excitatory and Type 2 inhibitory neurons are reciprocally connected, maintaining a precise E/I balance. Top-down feedback projections (orange) targeting Type 2 interneurons modulate the E/I balance. **(b)** Example synaptic currents into one excitatory neuron showing excitatory, Type 1 inhibitory and Type 2 inhibitory currents. **(c)** Illustration of E/I assembly dynamics and the effect of top-down feedback. In the absence of feedback, Type 2 inhibitory currents are proportional to the total feed-forward currents *I* ^ff^ *= I* ^exc^ *− I* ^inh1^ due to precise balance, allowing feed-forward currents to change neuronal firing rates while ensuring that the balance error Δ *=* 0 at all times. Top-down feedback into Type 2 neurons (orange) causes deviations from the balanced regime, leading to changes in firing rate and non-zero Δ. **(d)** Example time series of excitatory output firing rate (top), Type 2 inhibitory current (middle), and balance error Δ (bottom) for positive and negative feedback into Type 2 interneurons. Initially the neuronal input is precisely balanced resulting in Δ *≈* 0. Additional top-down input to Type 2 neurons suppresses firing while increasing inhibitory current resulting in a negative error encoding (Δ < 0). Conversely, Type 2 m ediated dis-inhibition results in a positive error encoding (Δ > 0). **(e)** Illustration of the BCP rule in the absence of top-down feedback. Regardless of feed-forward input strength (*I*^ff^), no plasticity is induced. **(f)** Schematic of the weight change 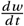 as a function of Δ. Sign and magnitude of synaptic plasticity are directly proportional to Δ. **(g)** BCP in a simulated patch-clamp paradigm (left), in which recurrent Type 2 inhibition is not driven by excitatory activity but is under experimental control. Synaptic weight change 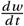 as a function of presynpatic stimulation for different values of Type 2 inhibition (right). In this case, BCP resembles a Hebbian learning rule in which the amount of inhibitory current controls the plasticity threshold.

We model excitatory neurons as leaky integrators with membrane potential *u*_*i*_ and rectified linear output firing rate *r*_*i*_ *=* (*u*_*i*_)_*+*_ of neuron *i*. For slowly changing input currents, the membrane potential is approximately

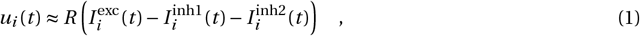

where *R* is the input resistance. We speak of neuron-specific “precise balance” [35] when the net feedforward currents are proportionally matched by recurrent Type 2 inhibition:

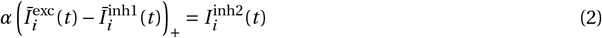

where *α <* 1 is a positive parameter and *Ī*^*x*^ denotes a short-term moving average over the respective synaptic current (Methods). We further define the balance error Δ_*i*_ of neuron *i* as follows:

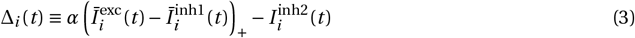

To test whether Δ could serve error encoding, we simulated an assembly of excitatory and inhibitory neurons in which the connectivity between Type 2 inhibitory neurons and excitatory neurons was optimized to ensure Δ_*i*_ *≈* 0 for any input (Fig. 2a,b; Methods). Moreover, Type 2 interneurons received top-down feedback, a common connectivity motif observed in neurobiology [63], causing changes in excitatory firing rates *and* deviations from precise balance reflected in non-zero Δ (Fig. 2c). Specifically, more Type 2 inhibition due to excitatory feedback suppressed excitatory firing rates and generated negative Δ (Fig. 2d) Conversely, Type 2 disinhibition caused increased excitatory firing rates and positive Δ. Thus Δ’s dynamics are consistent with a bidirectional error signal driven by top-down feedback to inhibitory interneurons.

To see whether the local errors Δ_*i*_ could serve as an error signal for error-driven learning, we have to make assumptions about the underlying learning algorithm. We now focus on the general class of local-error-correcting learning algorithms which includes backpropagation of error (backprop) and predictive-coding based learning algorithms as specific limiting cases [5, 6, 9, 11]. In accordance with the circuit introduced above (cf. Fig 2a), the required feedback is communicated through top-down control inputs to Type 2 interneurons. Given these constraints, the learning rule can be derived from an adaptive control theory framework (Methods). It is given by:

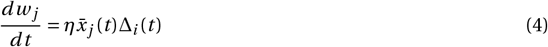

where *w*_*j*_ is the synaptic weight from presynaptic neuron *j, η* is a small positive learning rate and 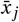 is a presynaptic activity trace. Because Δ_*i*_ corresponds to the balance error introduced above, we refer to Equation (4) as the balance-controlled plasticity (BCP) learning rule. In contrast to Hebbian learning rules, plasticity changes in BCP are independent of the total feed-forward drive into the neuron or the corresponding excitatory firing rate (Fig. 2e) as long as synaptic currents are precisely balanced. Instead, negative Δ *<* 0 results in long-term depression (LTD) while Δ *>* 0 triggers long-term potentiation (LTP) independent of the magnitude of postsynaptic firing rate (Fig. 2f). For feed-forward Type 1 inhibitory synapses, BCP operates with reversed polarity: negative Δ *<* 0 induces inhibitory LTP while positive Δ *>* 0 leads to inhibitory LTD. Importantly, for an individual neuron under tonic Type 2 inhibition, the BCP rule resembles a classic Hebbian plasticity rule whose plasticity threshold depends on Type 2 inhibition [56, 64] (Fig. 2g). Thus, BCP reconciles existing phenomenological plasticity models with the notion of error-driven synaptic plasticity through assembly-specific error signals.

### BCP supports plausible target-based learning through feedback control

Next we sought to understand whether BCP supports target-based learning [2, 9, 11]. To address this question, we designed a learning task in which assembly neurons needed to become selective to distinct input stimuli according to top-down feedback. Assembly neurons received temporally dynamic inputs from a population of 20 input neurons collectively encoding three continuous latent input signals (Fig. 3a,b; Methods). The connectivity between Type 2 inhibitory and excitatory neurons in the assembly was optimized to ensure precise E/I balance (Methods). We assessed the circuit’s selectivity in an open-loop setting (i.e. without top-down feedback) by presenting each stimulus in isolation. Before learning, assembly neurons maintained precise balance and showed nonselective responses to all stimuli (Fig. 3c). To test whether feedback could bias the assembly toward one of the input stimuli, we added feedback control to Type 2 interneurons proportional to a supervised target selectivity (Fig. 3b; Methods). In this closed-loop setting, feedback induced stimulus-specific balance perturbations when the response of the assembly did not align with the target, causing the assembly to respond strongly to a specific input signal and weakly to the other two. For instance, when we set A as the target, feedback caused disinhibition during stimulus A which encoded a positive error. Conversely, increasing Type 2 inhibition during non-target stimuli B and C encoded negative balance errors (Fig. 3c). As a result, the assembly’s selectivity and its associated open-loop response shifted toward A, while the required magnitude of the feedback control signal decreased monotonically (Fig. 3d). Importantly, this change in selectivity persisted in the absence of top-down feedback. To assess if these plasticity changes were reversible, we subsequently switched the output target to Stimulus B, causing a gradual loss of selectivity for Stimulus A and an acquired selective responses to B (Fig. 3d). These results show that top-down feedback can control plasticity through specific changes in E/I balance and enable target-based learning.

**Figure 3.**
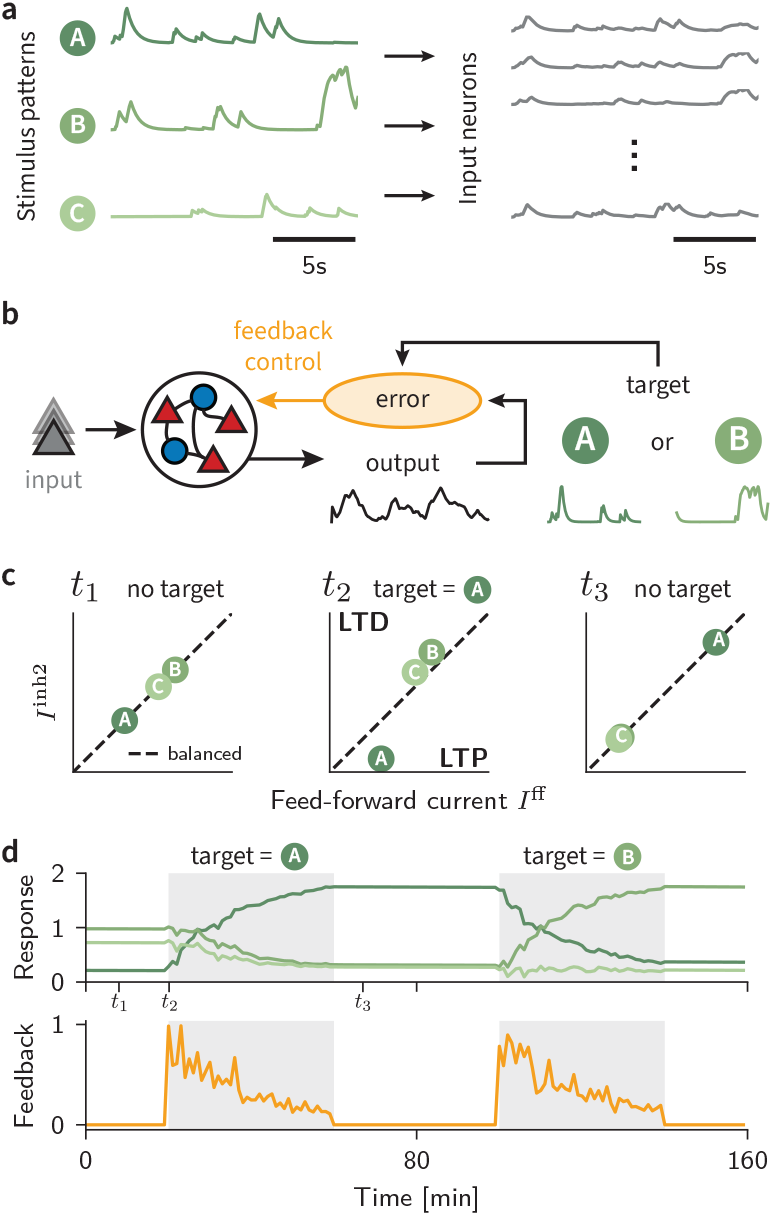
BCP supports plausible target-based learning through feedback control. **(a)** Schematic of the input paradigm. Three latent stimulus timeseries (A, B, C) are mixed linearly to yield 20 model inputs. **(b)** Sketch of the E/I assembly model. The assembly receives bipolar inputs through feedforward synapses subject to balance-controlled plasticity (BCP). The firing rate output of the excitatory assembly is compared to a given target signal. Deviations from the target result in a bidirectional error. Feedback control connections are set up such that their input to Type 2 interneurons reduces this error (Methods). **(c)** Feedforward input versus recurrent inhibition (Type 2) for the three different stimuli in isolation, evaluated at three timepoints of learning indicated in panel (d). Without a target signal (left), feedback control signals are zero and the inputs are precisely balanced (cf. Eq. (2)). When a target is given (middle), the balance is tipped in favor of LTP for the target pattern, whereas the two non-target patterns are suppressed. After learning (right), inputs are precisely balanced again, but the evoked feed-forward currents of each pattern changed due to synaptic plasticity. **(d)** Assembly response to the three target stimuli in the open-loop setting (top) and the strength of the feedback signal in the closed-loop setting (bottom) over time. Initially, there is no target and the response to all stimuli remains constant. When a target comes on (shaded regions), a feedback signal immediately corrects for any output error. Feedback signals decay slowly over time due to ongoing BCP which restructures the feedforward weights such that less feedback is required to produce the target output.

### BCP enables online learning in multi-layer networks

We now wondered whether top-down control of BCP could serve as a possible circuit mechanism capable of controlling the sign of plasticity in multi-layer networks. Since many learning algorithms require top-down feedback and local error signals controlling plasticity—irrespective of the specific objective function they optimize and which credit assignment strategy they use—we focused on supervised learning with deep feedback control [9]. Unless mentioned otherwise, we compute optimal top-down feedback signals from the network Jacobian (Methods), which requires full knowledge of the forward weights. Although this credit assignment strategy is considered biologically implausible, this choice allowed us to focus on the mechanism for encoding local error signals without having to commit to any specific biologically plausible approximation of backprop. With these thoughts in mind, we constructed a network model in which 20 periodic input units project to a hidden layer consisting of 240 excitatory and 60 Type 2 inhibitory neurons organized into 20 distinct E/I assemblies (Methods). The assembly output projected to two output neurons whose activity we interpreted as X-Y coordinates. The learning task was to map the periodic inputs to a predefined X-Y target trajectory corresponding to the outline of a turtle (Fig. 4a,b). The network output did not resemble the target prior to training (Fig. 4b). As before, we initialized synaptic weights to ensure E/I balance in the absence of feedback (Fig. 4c). We let input-to-hidden synapses evolve according to the BCP learning rule (cf. Eq. (4)), while output synapses learned according to the standard delta rule ([65]; Methods). In the closed-loop setup, Type 2 inhibitory neurons in the hidden layer received assembly-specific feedback control signals proportional to the output error that conveyed credit information. The resulting E/I balance deviations Δ collectively improved the match between the network output and the target trajectory (Fig. 4b,c), but showed only a weak correlation with neuronal firing rates on a single-neuron level (Fig. 4d,e). Because Δ_*i*_ determines the sign of plasticity in a neuron-specific manner, this behavior is different from classic Hebbian learning rules in which the sign of plasticity is usually correlated with postsynaptic firing rate [20–27] or by a global third-factor as in three-factor learning rules [66, 67].

**Figure 4.**
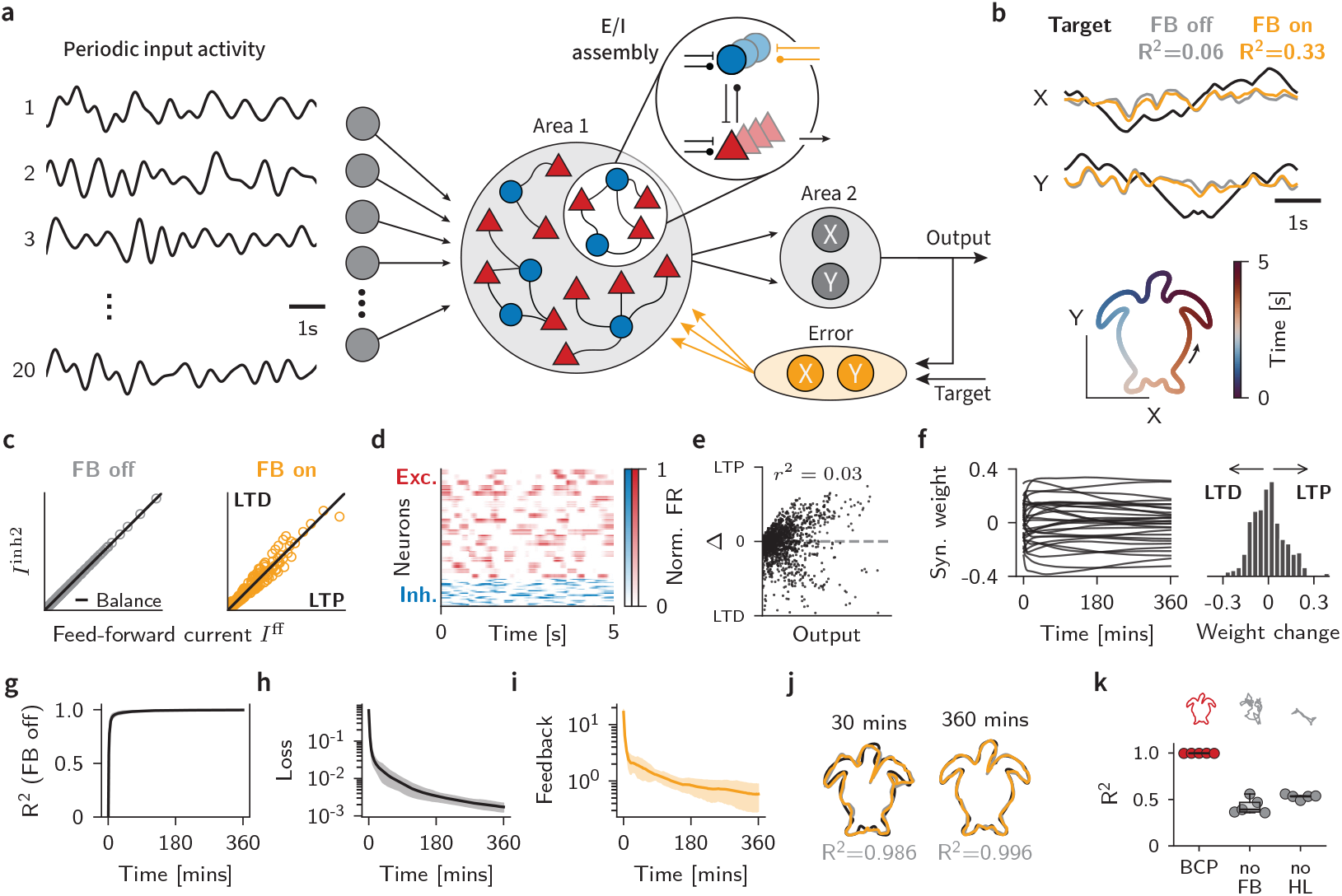
BCP enables online learning in multi-layer networks through feedback-guided plasticity. **(a)** Schematic of input stimulus and network architecture. 20 periodic inputs serve as inputs to a hidden layer organized into 20 E/I assemblies. Hidden layer neurons are connected to two output neurons encoding X-Y coordinates. Inhibitory neurons receive continuous feedback from a top-down controller proportional to the output error. **(b)** Target trajectory as a function of time (black) versus network output without (gray) and with top-down feedback (orange). Feedback nudges the output toward the target trajectory, resulting in increased *R*^2^. **(c)** EI balance for each neuron without (left) and with feedback (right). Each point corresponds to the synaptic currents into one excitatory neuron in the hidden layer averaged over 1s. Feedback induces bidirectional deviations from E/I balance that encode error signals. **(d)** Heat map of hidden-neuron firing rates over one trajectory before learning. **(e)** Synaptic weight changes as a function of postsynaptic firing rate. Each dot represents the postsynaptic firing rate and the corresponding neuron-specific Δ averaged over 1s. **(f)** Synaptic weight evolution of representative input-to-hidden weights (left). Distribution of weight final changes (right). **(g)** *R*^2^ during training. **(h)** Training loss over training (mean ± SD, n=5 networks). **(i)** Feedback magnitude as a function of time. **(j)** Output trajectory after 30 minutes and 6h respectively, demonstrating precise target matching. **(k)** Performance comparison (*R*^2^) across three conditions: BCP with feedback, no feedback (No FB), and no hidden layer (No HL). Control conditions fail to learn the task, showing that assembly specific feedback is necessary (mean ± SD, n=5 networks).

Next we switched on BCP and simulated the network for 360 minutes simulated time, corresponding to 4320 trajectory iterations, while periodically monitoring network performance in the open loop setting (Methods). BCP caused continuous synaptic weight changes in the hidden layer for which LTD and LTP were approximately balanced across synapses (Fig. 4f) and monotonically improved the match of the output to the target (Fig. 4g,h). At the end of the simulation, the network output trajectory closely matched the target(Fig. 4j) with an *R*^2^ *=* 0.996.

To test whether top-down feedback was necessary to solve the task, we repeated the same simulation without feedback, thereby restricting plasticity to output synapses. The resulting network failed to solve the task (*R*^2^ *=* 0.428; Fig. 4k). Similarly, when we removed the hidden layer entirely by directly mapping input nodes into the readout units, the network was unable to solve the task (*R*^2^ *=* 0.531). Lastly, to check whether the specificity of feedback connections was critical, we implemented random feedback projections [1, 68], either targeting entire assemblies or individual Type 2 interneurons, and found that networks could still successfully solve the task (Supplementary Fig. S2). These results demonstrate that BCP in combination with E/I assemblies and targeted feedback signals derived from an adaptive control theory framework supports learning in multi-layer networks.

### Learning is robust to moderate overlap in E/I assemblies

Thus far, we had considered idealized non-overlapping assemblies in the hidden layer in which each excitatory and each Type 2 inhibitory neuron belonged to a single assembly. However, it seems unlikely that such a clear assembly separation exists in neurobiology. Therefore, we sought to understand how robust the findings are to assemblies with overlap [69–71]. To address this question, we systematically varied the degree of overlap (Fig. 5a) by randomly assigning both excitatory and inhibitory neurons to additional assemblies (Methods). We characterize the degree of overlap as the ratio of additional assembly memberships to total assembly memberships.

**Figure 5.**
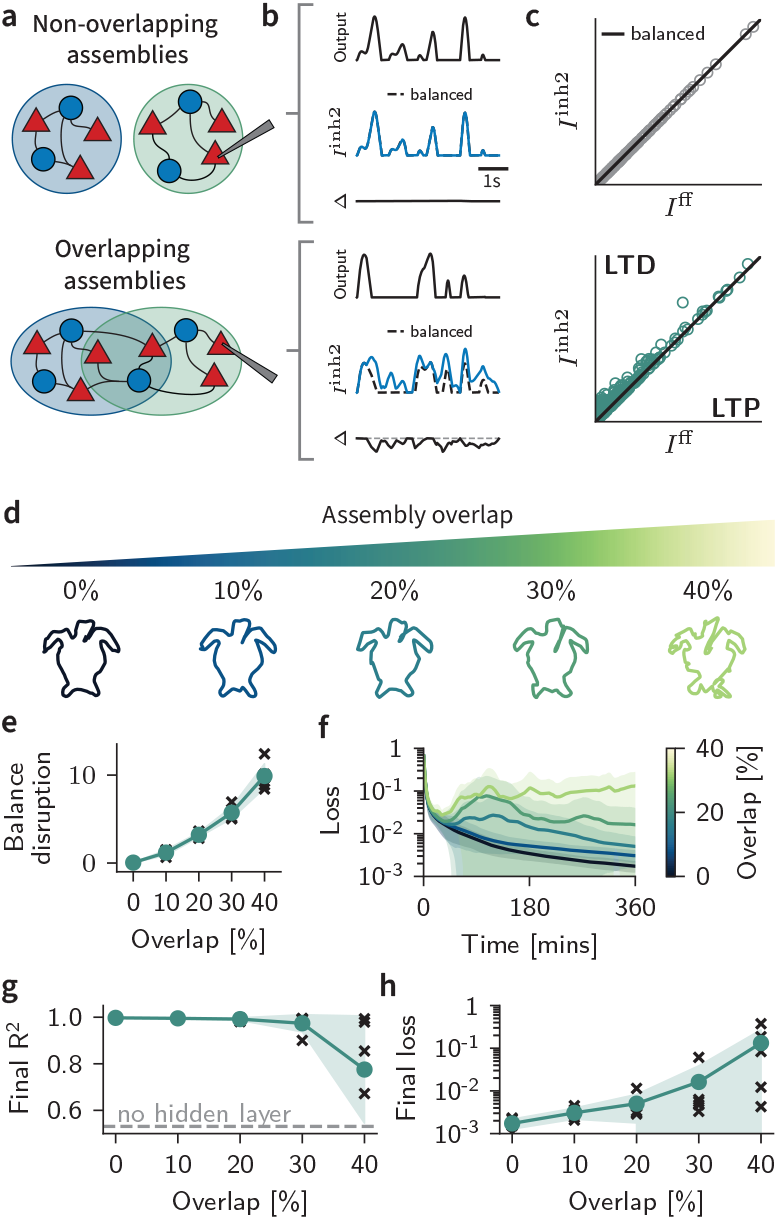
Learning is robust to moderate overlap in E/I assemblies. **(a)** Schematic illustration of non-overlapping (top) and overlapping (bottom) E/I assemblies. **(b)** Representative firing rate output, Type 2 inhibitory currents, and balance error Δ traces over time from non-overlapping (top) and overlapping (bottom) assemblies. Overlap induces deviations from the E/I balance target. **(c)** Scatter plot of E/I balance from the neuronal population in non-overlapping (top) and overlapping (bottom) networks. Each point represents average currents over one second for one hidden layer excitatory neuron. Overlap induces systematic deviations toward excess inhibition. **(d)** Output trajectories of networks with different levels of overlap trained on the trajectory matching task (see Fig.4). **(e)** Balance disruption, quan-tified as Σ_*i*_ |Δ_*i*_ |, as a function of overlap for individual networks (crosses) and mean (green; *n =* 5). **(f)** Temporal evolution of training loss for different degrees of over-lap. More overlap leads to unstable learning and worse performance. **(g)** Final R^2^ after 360 minutes of training on the trajectory matching task. **(h)** Same as (g) for the loss.

We first examined how assembly overlap affected E/I balance in hidden layer neurons. We found that assembly overlap induced a systematic disruption of the precise E/I balance, predominantly toward excess inhibition (Fig. 5b,c). This result can be understood because overlap causes excitatory neurons to receive input from and provide feedback to multiple inhibitory microcircuits. Consequently, the systematic disruptions induced by E/I assembly overlap might impair task performance when using E/I balance deviations as error signals for learning.

To test this hypothesis, we quantified the magnitude of balance disruptions for different degrees of assembly overlap (Fig. 5d; Methods). Doing so revealed that the degree of systematic balance disruption scaled monotonically with increasing assembly overlap as expected (Fig. 5e). We then trained these networks on the trajectory matching task as before (cf. Fig. 4) and measured final loss and *R*^2^. While all networks showed initial decreases in task loss, some configurations exceeding 10% overlap exhibited unstable learning dynamics (Fig. 5f). Notably, this effect had a strong initialization dependence as some networks with 40% overlap achieved comparable performance to non-overlapping architectures, while others showed severe impairment. Overall, we observed a graceful degradation of final performance with increasing assembly overlap (Fig. 5g,h). These results demonstrate that BCP learning is robust to moderate deviations from an ideal E/I assembly structure, whereas increasing levels of overlap degrades learning performance.

### BCP enables training of deep networks

So far we focused on a simple trajectory matching task in a network with a single hidden layer. To test whether BCP would also be conducive for learning for more complex classification tasks in multi-layer networks, we trained networks with either one or three hidden layers, each comprising 2048 excitatory and 512 inhibitory neurons organized into 128 assemblies (Fig. 6a), on Fashion MNIST, a ten-way classification task (Methods). In all networks, the output layer contained ten neurons corresponding to the ten classes. As before, we ensured assembly-specific feedback weights tuned to convey top-down feedback signals from a proportional controller at the network output (Methods). Given the static nature of Fashion MNIST inputs, we presented each input image until neuronal dynamics converged before updating synaptic weights based on the balance error Δ. We trained all networks for 50 epochs while monitoring performance on a held-out validation dataset and measured final performance on the test set.

**Figure 6.**
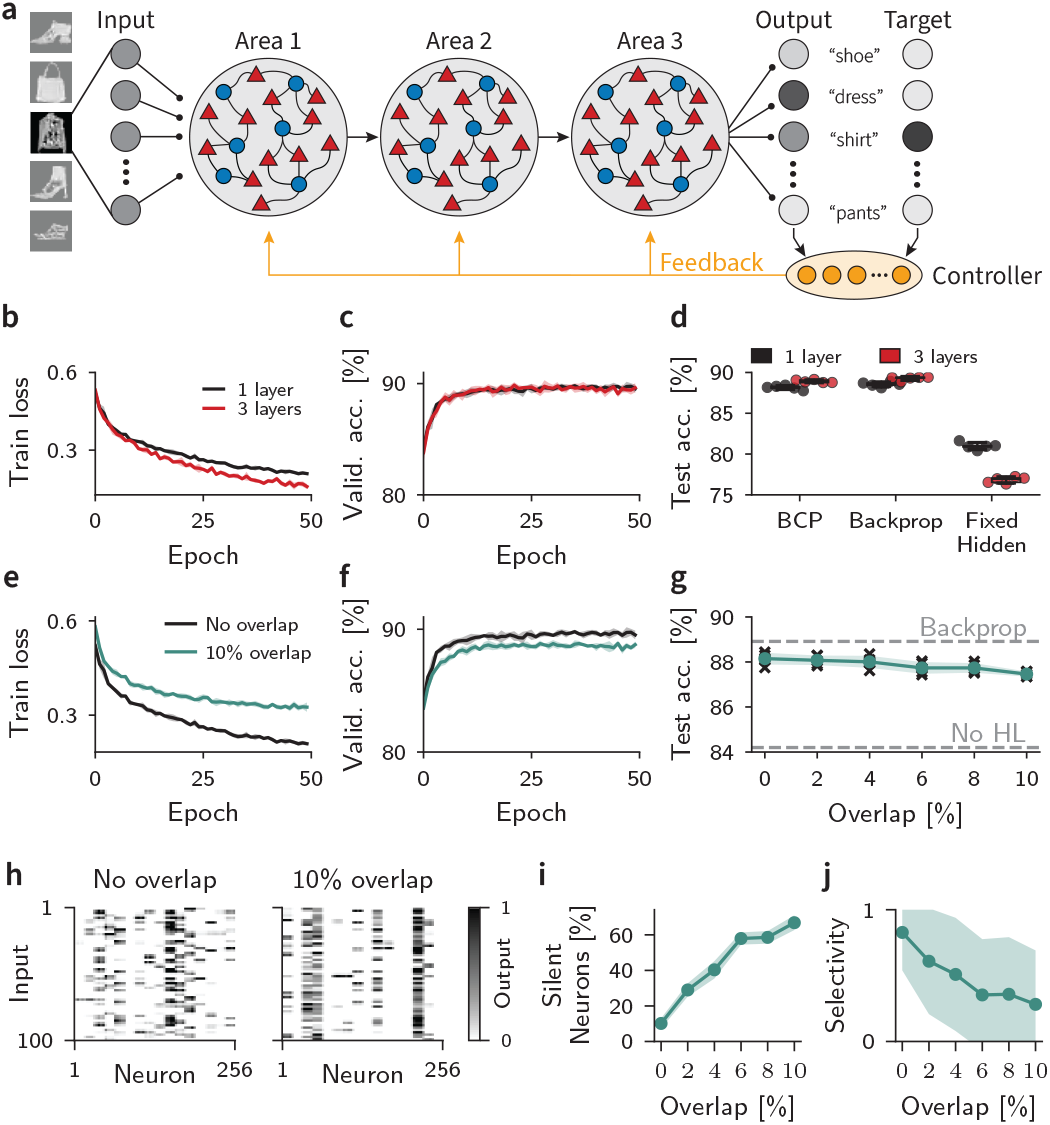
BCP enables training of deep networks on a computer vision benchmark. **(a)** Schematic of network for Fashion MNIST classification. Each hidden layer contains 2048 excitatory and 512 inhibitory neurons organized into 128 assemblies with assembly-specific feedback. **(b)** Evolution of training loss for networks with one or three hidden layers. **(c)** Evolution of validation accuracy during training. **(d)** Final test accuracy (*n =* 5). **(e**,**f)** Same as (b,c) for one hid-den layer with 0% and 10% assembly overlap. **(g)** Final test accuracy as a function of assembly overlap. **(h)** Post-training activity of 256 representative hidden layer excitatory neurons in a network without overlap (left) and with 10% overlap (right). **(i)** Percentage of silent neurons after training as a function of assembly overlap. **(j)** Neuronal selectivity measured as life-time sparseness post training as a function of assembly overlap. In all panels, shaded regions and error bars correspond to mean ± SD across *n =* 5 networks trained with different random seeds.

The training loss decreased during training of all networks (Fig. 6b) resulting in increasing classification accuracy (Fig. 6c). Networks with one hidden layer achieved a mean test accuracy of 88.1 *±* 0.3%, while networks with three hidden layers reached a mean test accuracy of 88.7 *±* 0.3%. For comparison we trained comparable networks using backpropagation which achieved comparable performance. In contrast, networks in which we trained the readout weights but not the hidden-layer weights performed substantially worse, indicating that feedback-driven plasticity in hidden layers was required for high task performance (Fig. 6d). These results demonstrate that BCP can successfully train deep networks to competitive performance levels on a standard machine learning benchmark.

To assess whether the resilience of BCP to assembly overlap extends to more complex learning scenarios, we trained single-hidden-layer networks with overlapping assemblies on the Fashion MNIST task. Networks with 10% overlap exhibited slower learning trajectories and converged to lower performance levels compared to non-overlapping networks (Fig. 6e,f). To quantify the relationship between assembly overlap and final performance, we trained networks with varying degrees of overlap (0-10%) and evaluated their test accuracy. We found that that final performance decreased slightly with increasing overlap percentage (Fig. 6g), consistent with our findings from the trajectory matching task.

To understand how assembly overlap affected network dynamics and learning performance, we analyzed the activity of hidden neurons in response to Fashion MNIST inputs after training. In networks with 10% overlap, we observed that several neurons failed to respond to any input, indicating that the network did not utilize these neurons for classification (Fig. 6h). We quantified the number of silent neurons and found that the percentage of these silent neurons increased with assembly overlap (Fig. 6i). While only 10.2 *±* 3.2% of neurons were silent in non-overlapping networks, this proportion rose to 66.8 *±* 4.1% in networks with 10% overlap.

To investigate how this increase in silent neurons affected the representation of task-relevant information in the network, we calculated the lifetime sparseness of hidden layer neurons across different inputs. This analysis revealed that the selectivity of the remaining active neurons decreased with assembly overlap (Fig. 6j). Together, these results suggest that assembly overlap impairs learning through two mechanisms: by reducing the number of neurons available for encoding task-relevant information and by decreasing the selectivity of the remaining active neurons.

### BCP captures signatures of *in-vivo* interneuron activity during learning

A plethora of experimental studies suggests that inhibitory circuits play essential roles in gating synaptic plasticity and learning [43, 45, 46, 48, 52, 53, 72–79]. More specifically, several studies implicated top-down activation of vasoactive intestinal peptide (VIP) interneurons with learning. We wanted to understand to what extend BCP could explain the results by Krabbe et al. [46] who showed that VIP interneuron activation in the Basolateral Amygdala (BLA) is required for learning in an auditory fear conditioning experiment. In this experiment, freely moving mice were exposed to ten conditioned stimulus (CS)-unconditioned stimulus (US) pairings over the course of two days while the activity of VIP interneurons was assessed via two-photon Ca2+ imaging through a head-fixed miniature microscope (Fig. 7a). The authors found that VIP interneurons responded strongly to the US in naive animals and that this response reduced over the course of ten pairings, accompanying the acquisition of a behavioral freezing response to the CS (Fig. 7b). To test whether BCP could account for these findings, we used it to train E/I assembly networks on the same task (Fig. 7c). In the model, the CS corresponded to sparse noisy input and the network was trained to output the freezing probability (Methods). We modeled the US as a negative valence target for the network output and implemented proportional error feedback through a dis-inhibitory circuit motif mediated by Type 2 interneurons (Methods). We simulated ten CS-US pairings and recorded the amount of US-driven dis-inhibitory feedback as well as freezing responses as in the experiment. The model output acquired a reliable freezing response which was accompanied by a reduction in US-driven dis-inhibitory feedback (Fig. 7d) similar to the experiment. In the experiment, optogenetically inhibiting the US response of VIP interneurons impaired learning (Fig. 7e). To test whether dis-inhibitory feedback is also required for learning in the model, we inhibited the VIP population by reducing the magnitude of the control signal by a factor of ten. This manipulation led to a stark reduction in freezing probability similar to the experimental data (Fig. 7f). Thus, our model can explain VIP interneuron activity during learning, suggesting a putative role for these inhibitory microcircuits in encoding top-down error signals in the BLA.

**Figure 7.**
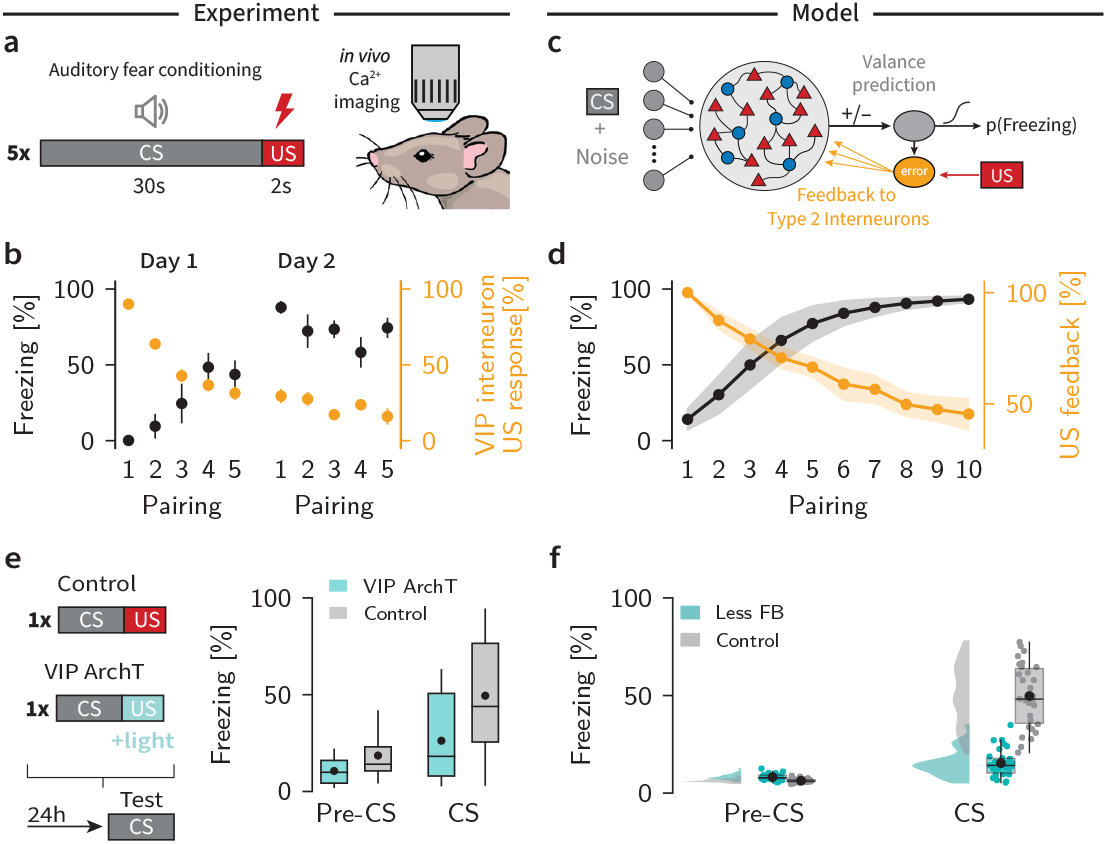
BCP captures signatures of fear learning in the rodent BLA. **(a)** Schematic of the fear conditioning experiment by Krabbe et al. [46]. Freely moving mice were exposed to five pairings of CS (tone) and US (footshock) while the activity of VIP interneurons in the BLA was recorded using a genetically encoded Ca2+ indicator. Learning was assessed by recording the animal’s freezing response to the CS presentation. **(b)** Time course of freezing responses and VIP neuron activation over sessions in the experiment [46]. Mice gradually acquired a freezing response to the CS over 10 pairings over 2 days (black). Learning was accompanied by decreasing VIP interneuron responses to the US (orange). Data extracted from [46]. **(c)** Schematic of the BLA network model trained with BCP. The output of the network serves as a valance prediction, which is translated through a sigmoidal activation function into freezing probability. Error signals are computed by comparing valence prediction to a target value associated with the US and fed back to the network through a dis-inhibitory circuit (orange; Methods). **(d)** Timecourse of freezing and VIP feedback during fear learning in the model. Learning corresponds to increasing feezing probaility (black) accompanied by decreasing US-driven dis-inhibitory feedback (orange)mirroring key elements of the experiment (cf. (b)). Shaded areas correspond to the standard deviation (*n =* 30 simulation runs). **(e)** Left: Schematic of the experiment [46] in which the response of VIP interneurons to the US was optogenetically decreased during CS-US pairing (VIP ArchT condition). Right: Freezing behavior before and after fear conditioning indicates that US-driven VIP interneuron activation during pairing is necessary for learning. Data extracted from [46]. **(f)** Freezing probability in the model before (left) and following learning (right) for simulted ArchT and the control condition. Reducing the amount of US-driven feedback in the model leads to impaired learning, which is consistent with the experimental findings (cf. (e)).

To understand whether BCP could also capture interneuron dynamics in the cortex during learning, we trained a multi-layer network model with BCP on a motor learning task (Fig. 8a) known to require cholinergic top-down feedback to VIP-somatostatin (SOM) dis-inhibitory microcircuits in primary motor cortex (M1) [47]. Networks learned to generate a motor output resembling a lever push in response to an auditory cue over the course of 2000 training iterations (Fig. 8b,c), as in the experiment by Ren et al. [47] (Fig. 8d). Importantly, Type 2 neurons in the model showed bi-directional activity changes over learning, with an overall trend towards increased activity (Fig. 8e), consistent with activity changes in SOM interneurons in the experiment (Fig. 8f). In contrast, the amount of top-down feedback to almost all E/I assemblies in the model decreased over learning (Fig. 8g), closely resembling the experimental time course of VIP interneuron activity in the experiment (Fig. 8h). Thus, the evolution of activity of Type 2 interneurons and top-down feedback during learning in the BCP framework captures the activity changes observed in SOM and VIP interneurons in M1, respectively, suggesting a role for these inhibitory microcircuits in local error encoding.

**Figure 8.**
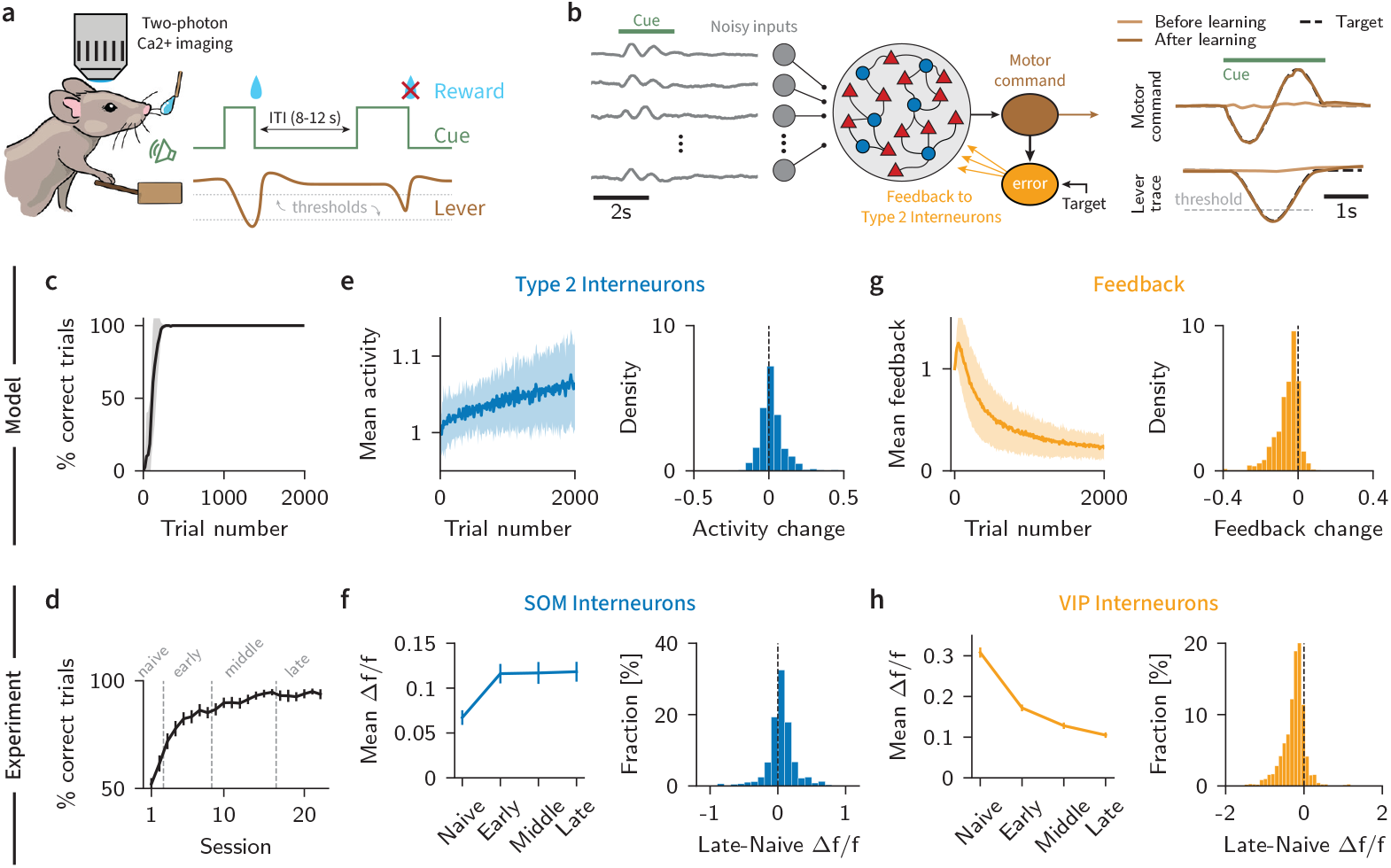
BCP captures activity changes of M1 interneurons during motor learning. **(a)** Schematic of the motor learning experiment by Ren et al. [47]. Mice were trained to push and pull a lever in response to an auditory cue to obtain a water reward over 22 sessions spread across multiple days. In all sessions, the activity of VIP and SOM interneurons in M1 was recorded using two-photon Ca2+ imaging. **(b)** Schematic of the M1 network model trained with BCP. In each trial, the model receives noisy cue-driven inputs that are processed by a hidden layer with *n =* 40 E/I Assemblies. A single output neuron was trained to follow the simulated time course of a push-pull movement in response to the cue. **(c)** Evolution of network performance during task learning. The shaded region indicates one standard deviation (*n =* 10 networks). **(d)** Performance of mice across training sessions. Data extracted from [47]. **(e)** Left: Evolution of mean Type 2 interneuron activity in the model during motor output across trials. Right: Distribution of learning-induced Type 2 interneuron activity changes in the model. **(f)** Left: Mean activity of SOM interneurons in M1 during movement at different stages of learning. Right: Distribution of activity changes across SOM interneurons in M1. Data extracted from [47]. **(g)** As in panel **(d)**, displaying the magnitude of top-down feedback in the network during motor output. **(h)** As in panel **(e)**, for VIP interneurons. Data extracted from [47].

## Discussion

In this article, we investigated a circuit-level mechanism for error-driven learning based on deviations from precise E/I balance. We introduced balance-controlled plasticity (BCP), a plasticity model that assumes that neurons and synapses can detect both the amount and source of inhibitory input in a cell-type-specific manner. We showed that BCP supports online learning in multi-layer networks when suitable top-down feedback influences local microcircuits. Finally, models in which feedback targets an inhibitory subpopulation naturally replicate *in-vivo* activity changes of SOM and VIP interneurons during learning [46, 47].

Neuron-specific errors that influence plasticity are essential for many plasticity algorithms—most notably the ones solving the spatial credit assignment problem. How the brain accomplishes such error specificity remains an open question [2, 13]. Recent models proposed mechanisms such as separate temporal phases [14] for feed-forward vs error encoding or argued that electrotonically segregated dendrites are ideally suited for integrating error or target signals [7, 9, 11, 80–83]. However, such a separation of feed-forward activity and error encoding is often ad-hoc, not always easy to reconcile with neuronal data, and raises conceptual questions. On important question is how error signals can be sensed at synapses targeting the soma or basal dendrites if they are encoded in the apical tuft dendrite? While some models compellingly argue that an intricate burst multiplexing code based on nonlinear interactions of apical and basal dendritic compartments constitute a plausible mechanism that circumvents this problem [15, 84], these models chiefly apply to Layer 5 pyramidal cells, which possess the required well-separated apical and basal dendritic compartments and exhibit bursting activity consistent with the notion of multiplexing [16]. However, this reliance leaves open the question of how other neurons, for instance in cortical Layers 2/3 or in subcortical brain areas, that do not meet these criteria, accomplish credit assignment.

In this article we have proposed one possible alternative. Since E/I currents are often precisely balanced, i.e., correlated, this create a null-space in which neuronal activity does not change. BCP’s core idea is to use this null-space for error coding. Consequently, BCP does not rely on any anatomical specialization or compartmentalization. Instead, it requires mechanisms to retain and break precise E/I balance in an orchestrated manner. While some of the models discussed above also posit a form of balance in their apical dendrite [7, 15, 84], our work shows that we can dispense with this dendritic compartment and effectively externalize it into a local recurrent microcircuit, i.e., an E/I assembly. What we gain through this seemingly small change is that the resulting learning rules are more closely aligned with phenomenological voltage-dependent plasticity models. A direct consequence is that E/I assemblies effectively replace individual neurons as the computational units in this model.

To enable learning in multi-layer networks, BCP needs to be accompanied by a credit assignment strategy. Here, our work draws from recent models that use strong top-down feedback to influence neuronal activity and plasticity [5, 11, 81, 82, 85–87]. These models can be derived from a minimization of control principle in which the error signal emerges from the difference between *open-loop* (in the absence of top-down feedback) and *closed-loop* activity (with feedback). A central challenge for these models remains that there is little evidence for explicit open and closed-loop phases in neurobiology. Previous models solved this issue by, once more, asserting segregated dendrites [5, 81, 82, 87] or dedicated error neurons [11] based on similar mathematical ideas [10]. Other recent work demonstrated that the need for separate phases or compartmentalization can be relaxed when temporally delayed top-down feedback induces firing rate changes detectable by an spike-timing-dependent plasticity (STDP) rule [88]. In contrast, BCP allows reformulating the problem by mapping the open-versus closed-loop contributions onto a distinct coding axis related to deviations from E/I balance. Here, the expected inhibitory current under precise balance encodes the *open-loop* activity, while the actual inhibitory input reflects the *closed-loop* state. This separation gives each neuron implicit access to its own *open-loop* activity without requiring additional circuitry or compartmentalization.

Importantly, mapping error signals to E/I balance instead of putative anatomical compartments also results in learning rules that more closely align with phenomenological plasticity models. We showed that BCP resembles a Hebbian plasticity rule with a dynamic plasticity threshold controlled by inhibition (cf. Fig. 2h). While plasticity thresholds are central to classic models of Hebbian learning [89, 90] and phenomenological plasticity rules [23, 24, 26, 91, 92], most learning rules do not depend explicitly on inhibition. Inhibition-dependent learning has been investigated in recent theoretical studies, which implicated it in receptive field formation and stabilization of learning [55–57, 64]. While these studies shared BCP’s core principles of E/I balance and an explicit inhibition dependence of plasticity at excitatory synapses, they did not study the implications of implicit error encoding for credit assignment in multi-layer networks.

Finally, our work suggests a reinterpretation of the basic computational units in biological neural networks. Specifically, E/I assemblies are the computational units in our model rather than individual neurons. This interpretation opens up exciting parallels to recent models on efficient coding [36, 93], low-rank networks [94, 95], and work showing that low-dimensional error signals are often sufficient for learning complex functions in deep networks [96]. Because the number of E/I assemblies puts an upper bound on the effective dimensionality of the neuronal manifold, we expect that brain areas encoding high-dimensional information comprise many small assemblies rather than a few large assemblies. Interestingly, there is indirect evidence for such E/I assembly structures, for instance, in the area Dp, the homologue of the olfactory cortex in zebrafish [37, 97].

### Putative bio-chemical mechanisms of BCP

Our model is built on the assumption that neurons can distinguish between excitatory and specific types of inhibitory input. Fundamentally, BCP’s balance condition (cf. Eq. (2)) can be interpreted more broadly as a balance between LTP-inducing and inhibition-controlled LTD-inducing plasticity mechanisms. While the concrete mechanism is unknown, several molecular pathways could be involved in mediating such balance control. First, inhibitory currents can precisely regulate local dendritic voltage [98], thereby controlling Ca^2+^ influx through NMDA channels [99] known to modulate plasticity [100]. Additionally, inhibition can directly modulate Ca^2+^ permeability of both NMDA receptors and voltage-gated Ca^2+^ channels through non-ionotropic GABA-B receptor signaling [101–103]. Another conceivable pathway extending beyond calcium signaling may rely on inhibitory inputs regulating local kinases and phosphatases involved in synaptic plasticity [104, 105]. While the model formulation in this article assumes these processes operate at synaptic timescales, biological plasticity mechanisms likely incorporate multiple timescales of balanced processes [106, 107]. Thus, the effects of E/I balance on plasticity may manifest more slowly than synaptic currents, suggesting that plasticity-relevant processes could still cancel even without millisecond-precision of the E/I balance [35].

### Limitations

We made several simplifying assumptions. First, we assumed that top-down feedback weights were optimized to deliver relevant credit signals to their target assemblies (except in Supplementary Figure S2, where we used random feedback). The need for appropriate feedback signals is a fundamental challenge shared by all biologically plausible credit assignment strategies [2, 13]. Still several studies have shown that this limitation can be mitigated by learning the feedback weights [9, 108–113] or by redesigning feedback pathways [114]. In this article, we did not focus on any particular candidate feedback mechanism. Instead we presented an effective error encoding mechanism which could work with most, if not all, biologically plausible credit assignment models that traditionally rely on separate compartments for receiving such error signals.

In this article we focused on feedback to Type 2 interneurons which provided recurrent precisely balanced inhibition. This choice was inspired by the prominent dis-inhibitory circuit motifs found in many brain areas [115] and involved in gating of plasticity [116]. However, it is important to note that our adaptive control theory framework is agnostic to how E/I balance is established and how it is influenced through feedback. Hence other circuit models and feedback pathways are possible. For example, BCP would yield a similar workable model in which feed-forward inhibition is responsible for precise balance and targeted by feedback (cf. Fig. S1). Similarly, feedback pathways relying entirely on excitatory neurons are possible too. As long as E/I balance is modulated in an orchestrated way, we expect BCP to work.

Another limitation of our work is that while our model requires E/I assemblies, it remains agnostic about how they are established and maintained during learning. In this article, we artificially enforced E/I assemblies for simplicity. In neurobiology, the required assembly connectivity would necessarily have to be self-organized during development and learning. Fortunately, several studies have shown how inhibitory plasticity can establish co-tuned E/I assemblies through decorrelation of excitatory activity [59–61, 117–121]. Still, maintaining E/I balance while using balance errors for learning poses a chicken-egg problem. Solving this problem may ultimately require cycling between different plasticity states. A promising candidate could be our wake-sleep cycle. For instance, balance errors would serve error-driven learning during wakefulness. In contrast, one of the roles of sleep may be to re-establish E/I balance [122], a notion that also seems consistent with work showing that inhibitory engrams emerge over prolonged periods *after* memory storage [123]. Beyond maintaining E/I balance, inhibitory plasticity could serve complementary functions in learning that we did not consider in our model. While we have considered circuits in which top-down feedback transiently modulates inhibitory activity to induce plasticity, other computational work explored networks in which top-down feedback drives cell-type specific inhibitory plasticity [124] that could allow for prolonged control over excitatory plasticity even after top-down feedback subsides.

Moreover, BCP is a simplified plasticity model which does not yield classic plasticity curves associated with low postsynaptic activity result in LTD whereas high activity levels yield LTP in the absence of inhibition. This choice was intentional for ease of analysis, but is not a fundamental limitation of the model. Importantly, we have shown previously that learning rules with an explicit inhibition dependence in which error signals are decoded through a linear approximation of the inverse of the neuronal transfer function [125] naturally recover classic plasticity curves while also providing rapid compensatory control of Hebbian plasticity through recurrent inhibition alone [126, 127].

Finally, our model does not *require* electrically well-separated dendritic compartments to encode error signals. However, beyond any doubt dendritic compartmentalization fulfills important computational functions [128–130]. Curiously, recent experiments have shown that E/I balance is maintained at the level of local dendritic segments [131], and that some interneurons form synapses on individual dendritic spines [98]. While we have only considered simple neuron models without dendritic compartments in this article, in future work we would like to extend the BCP framework to compartment-specific error signals through precise E/I balance that is maintained at the level of individual dendrites and explore whether such compartment-specific error encoding offers computational advantages beyond neuron-wide error signals.

### Experimental predictions

Our study makes several testable predictions. First, the BCP learning rule predicts that inhibitory interneuron activity should be modulated by top-down feedback during learning and thereby effect both excitatory activity *and* plasticity. While there is ample evidence for inhibitory gating of plasticity both *in vitro* [39–45, 53] and *in vivo* [46–49, 52, 54, 76, 77], most experiments did not separately control inhibition and neuronal firing rates because they are typically anti-correlated. To isolate balance effects from firing rate changes, we propose a closed-loop experimental learning paradigm. In this setup, stimulus strength or excitatory neuronal input is dynamically adjusted to maintain constant target firing rates in an excitatory population. At the same time recurrent inhibitory activity is modulated, e.g., optoge-netically, to break E/I balance. Under these conditions, BCP predicts observable plasticity at input synapses despite stable output firing rates. In addition, the direction and magnitude of plasticity should be a function of inhibitory activity.

Second, our model posits a dual role for inhibition, suggesting that inhibitory plasticity rules should differ based on the presynaptic population’s circuit function. In our model, we distinguish Type 1 inhibition, which acts as the negative part of a neuron’s receptive field, and Type 2 inhibition, which establishes precise balance. BCP predicts that Type 2 (primarily recurrent) inhibitory synapses should exhibit co-dependent plasticity to maintain balance [57], while the sign of plasticity at Type 1 (primarily feed-forward) inhibitory synapses should be governed by deviations from E/I balance. This prediction aligns with the diverse forms of inhibitory STDP observed experimentally [120, 132]. Importantly, the relationship between connectivity motifs and circuit function does not have to be rigid—some circuits may employ feed-forward inhibition to maintain balance [25, 133–135] while using lateral inhibition between different assemblies to shape receptive fields (c.f. Supplementary Fig. S1).

Finally, our model assumes that specific inhibitory interneuron subpopulations are modulated by top-down feedback during learning consistent with feedback control. This assumption is also a prediction. Specifically, we expect that such inhibition-mediated feedback responding to external stimuli early in learning can predict both behavioral and neuronal responses that emerge later in response to the same stimuli. Several studies have presented evidence for such (dis-)inhibitory feedback control [46, 49, 74, 78, 136] that guides learning and population activity. However, whether this top-down control predicts plasticity at the single neuron level remains unresolved.

In summary, our work introduces a biologically grounded mechanism by which neurons can encode local error signals through deviations from precise excitation-inhibition balance. This balance-controlled plasticity enables online credit assignment without requiring specialized compartments or phases, unifying normative principles with observed cortical microcircuit dynamics. By linking synaptic plasticity to transient deviations in E/I balance, our model offers a testable framework for how neuronal circuits may solve complex learning tasks using local information.

## Acknowledgements

We thank all members of the Zenke Lab for comments and discussions throughout this project. Thanks to Peter Buttaroni for providing helpful illustrations for figures in this manuscript. This project was supported by the Swiss National Science Foundation (Grant Number PCEFP3_202981), EU’s Horizon Europe Research and Innovation Programme (Grant Agreement No. 101070374, CONVOLVE) funded through SERI (Ref. 1131– 52302), and the Novartis Research Foundation.

## Author contributions

Julian Rossbroich: Formal analysis, Investigation, Methodology, Software, Visualization, Writing – original draft, Writing – review and editing; Friedemann Zenke: Conceptualization, Funding acquisition, Methodology, Supervision, Writing – original draft, Writing – review and editing.

## Data and code availability

The code to reproduce all modeling results reported in this article is available at https://github.com/fmi-basel/balance-controlled-plasticity

## Supplemental information

Supplementary Material document. Figures S1 & S2, Tables S1-S5 and supplementary methods.

## Methods

### Neuronal dynamics

Throughout this article, we simulated networks of *N*_E_ excitatory and 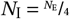 Type 2 inhibitory neurons with reciprocal connections. We write 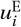 for the membrane potential of excitatory neuron *i* and 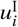 for the *i*-th inhibitory neuron. These membrane potentials evolved according to the following differential equations

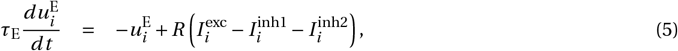

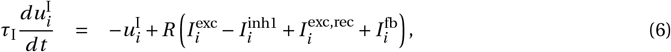

where 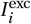 and 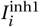 denote the excitatory and Type 1 inhibitory currents originating from external input populations, 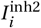 are Type 2 inhibitory currents, 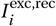 are recurrent excitatory currents and 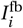 are top-down feedback currents to Type 2 inhibitory interneurons that can be either excitatory or inhibitory. Finally,*τ*_E_ and *τ*_I_ are the membrane time constants for excitatory and inhibitory neurons, respectively. For ease of notation we modeled synaptic currents in natural units assuming an input resistance *R =* 1.

#### Synaptic connections

Each input current was modeled as a weighted sum over presynaptic activities. In general, we denote the synaptic weights from presynaptic neuron *j* in population *A* to postsynaptic neuron *i* in population *B* as 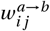. Neurons received excitatory and Type 1 (feed-forward) inhibitory inputs from a population of *N*_X_ neurons with activities 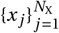. For ease of analysis, we assume instantaneous Type 1 feed-forward inhibition. In other words, we treated Type 1 inhibitory neurons as part of the input population and decomposed the input weights into excitatory and inhibitory components

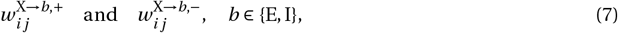

to denote the excitatory and inhibitory weights, respectively. Accordingly, the dynamics become

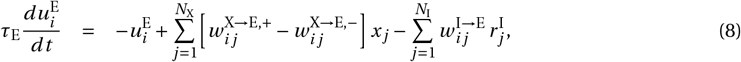

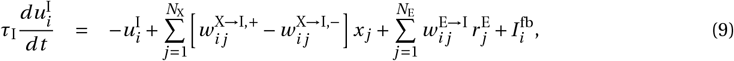

where 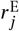 and 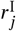 denote instantaneous firing rates that were computed as

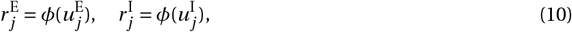

with the rectifying nonlinearity *φ*(*u*) *=* max{0, *u*}. To simplify further analysis, we combine the positive and negative feedforward weights into a single weight matrix as follows:

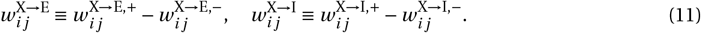

With these definitions, the dynamics of the E/I assembly populations simplify to

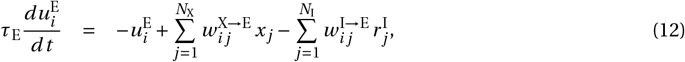

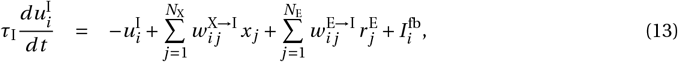

or, in vectorized notation,

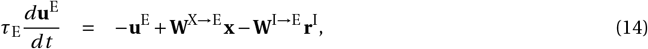

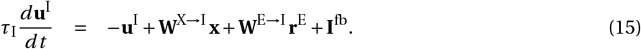

#### Readout neurons

Unless otherwise stated, we modeled the output of the network as a group of *N*_out_ linear readout neurons with the following dynamics:

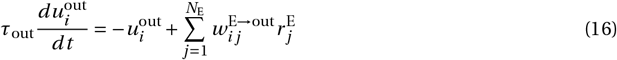

where *τ*_out_ is the readout time constant and number of readout neurons *N*_out_ depends on the specific task (see below).

### E/I Assembly structure

Neuronal E/I assemblies are the basic computational element in our model. The population of excitatory and Type 2 inhibitory neurons was divided into *N*_A_ assemblies, each containing 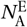 excitatory neurons and 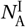 inhibitory neurons. The membership of neurons to these assemblies was specified by the membership matrices **M**^E^ and **M**^I^. In particular, for both membership matrices, the element

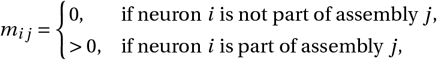

with the positive value quantifying the relative contribution of neuron *i* to assembly *j*.

#### Within-assembly excitatory synaptic weights

A neuron’s assembly memberships determine its recurrent connections within the network. We considered fixed E *→* I weights so that the recurrent weight from excitatory neuron *j* to Type 2 inhibitory neuron *i* is given by

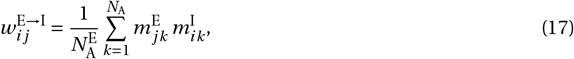

where 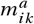 denotes the membership of neuron *i* in population *a* ∈ {E, I} to assembly *k*. This led to a connectivity structure,in which each inhibitory neuron receives recurrent input from excitatory neurons belonging to the same assembly, with the connection strength determined by the degree of membership of both neurons in that assembly.

#### Parameterization of feed-forward synaptic weights

Our model requires co-tuned E/I assemblies which we assumed as given, for instance, as the outcome of developmental or learning processes [59, 60, 62, 118]. Specifically, we assumed that neurons belonging to the same assembly share the same feed-forward tuning. Consequently, feed-forward weights to assemblies were the learnable parameters

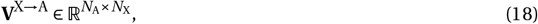

which map inputs to assembly space, in conjunction with the respective membership matrices. Specifically, this parameterization is expressed as

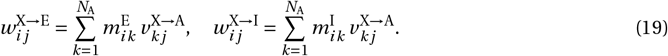

Similarly, we assumed that network readout neurons receive assembly-specific connections given by the learnable parameters

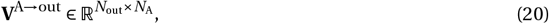

that determine the readout weights

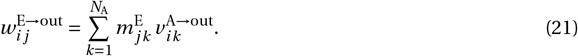

#### Initialization of assembly structure

In simulations with a single assembly (Figs. 2 and 3) or networks with a non-overlapping assembly structure (Figs. 4, 6, 7, 8), each neuron was assigned exclusively to one assembly. To that end, we partitioned the excitatory and inhibitory populations into *N*_A_ assemblies such that the numbers of excitatory and inhibitory neurons per assembly are given by

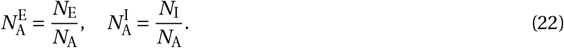

For each population *a ∈* {E, I}, with *N*_*a*_ denoting the number of neurons in population *a*, we defined a binary membership matrix

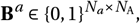

where

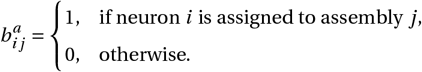

In the non-overlapping case, each neuron was a member of exactly one assembly. To account for heterogeneous contributions within an assembly, we created a weight matrix

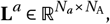

with entries independently sampled from a lognormal distribution with parameters *µ*_memb_ and *σ*_memb_. The final membership matrix was then defined via the elementwise product,

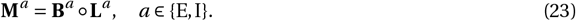

#### Initialization of overlapping assemblies

In networks with overlapping assemblies (Figs. 5 and 6), we started from the binary matrices **B**^*a*^ defined above for *a* ∈ {E, I}. We then introduced additional memberships by randomly setting extra entries to 1, yielding an augmented binary matrix 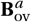. Since neurons may now have multiple memberships, we normalized each row of 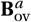 to ensure that the total membership weight per neuron equals one:

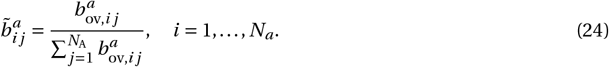

As before, we generated a weight matrix

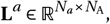

with lognormally distributed entries and defined the final membership matrix as

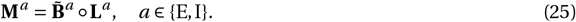

#### Quantification of assembly overlap

To quantify the degree of assembly overlap, we considered the ratio of additional entries in the augmented binary matrices 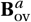 relative to the total number of entries in these matrices. In particular, we define the overlap ratio as

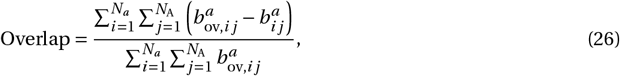

which ranges from 0 (no overlap) to values approaching 1 (very high overlap). In the main text, the overlap ratio is expressed as a percentage.

#### Optimization of Type 2 inhibitory connections for E/I balance

Throughout this paper, we required that E/I assemblies operate in a precise E/I balance regime (cf. Eq. (2)), in which Type 2 inhibitory currents maintain a proportional relationship with the net feed-forward activity. In contrast to previous computational studies that investigated how E/I balance can be established and homeostatically maintained through inhibitory plasticity mechanisms [25, 56, 57, 59], we chose a mechanism-agnostic approach in this article. To that end, we directly optimized the Type 2 inhibitory I *→* E synaptic weights to satisfy the balance condition in equation (2) according to

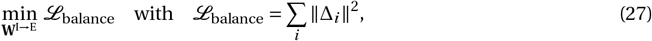

using standard gradient descent methods (see Supplementary Material Section 1.1). Minimizing the balance error Δ for each neuron at the time of network initialization ensured precise balance regardless of the network inputs that persisted during learning without the need for continuous optimization.

### Top-down error feedback

In our model, top-down feedback to Type 2 interneurons provides an adaptive control input that changes activity to minimize a loss at the network output [5, 81, 87]. We modeled the top-down currents **I**^fb^ as feedback control currents from a controller population **c** through excitatory or inhibitory feedback afferents with weights **Q**. Importantly, our framework makes little assumptions about the actual dynamics of the controller, its implementation, and the nature of the feedback weights, as long as the control currents lead to reducing the loss function (see also [9, 81]). In this article, we considered simple leaky proportional-integral controllers that follow the dynamics

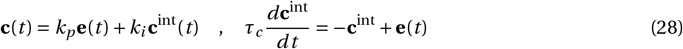

where 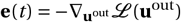 is the magnitude of the error at the network output, *k*_*p*_ is the proportional gain, *k*_*i*_ is the integral gain, and *τ*_*c*_ is the time constant of the integral controller.

#### Feedback weights

The feedback weights **Q** describe the effect of the controller on the inhibitory interneurons. In this article, we only considered direct linear feedback from the controller to the inhibitory interneurons, which could be interpreted as direct feedback projections from another brain area. However, other forms of feedback involving local mechanisms or non-linear pathways are possible [81]. Following previous work [81, 87] we chose a direct linear feedback mapping that approximates the network Jacobian (see Supplementary Material Section 1.2), such that

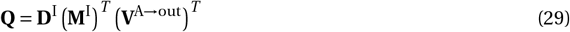

where **D**^I^ is a diagonal matrix containing the derivative of the rectified nonlinearity for each inhibitory neu-ron so that 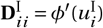. The inclusion of this derivative term ensured that top-down feedback only targeted active inhibitory neurons that operate in the linear regime.

### Balance controlled synaptic plasticity

Given the E/I assembly structure and top-down feedback connectivity discussed above, the BCP learning rule can be derived from a minimization of control objective function (Supplementary Material, Section 1.3). In brief, the idea is that learning should reduce the amount of control the controller has to exert to minimize the output error. Given linearized Type 2 inhibitory dynamics, we obtain the BCP learning rule for feed-forward weights **V**^X*→*A^ as the negative gradient of the least control loss function ℋ*=* ||**W**^I*→*E^**Qc**||^2^:

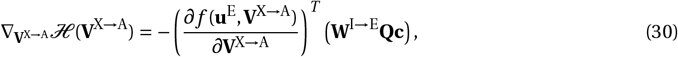

which yields the following gradient-based update rule for the feed-forward parameters **V**^X*→*A^:

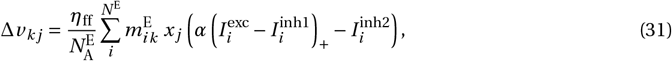

#### Online learning

Importantly, the derivation from a least control objective assumes that neuronal dynamics have converged to an equilibrium state for a given input *x*_*j*_. While this is possible for static inputs (Fig. 6), during online learning with dynamic input stimuli (Figs. 3, 4, 5, 7 and 8), this condition is not formally met. That is, the temporal dynamics of Type 2 inhibitory neurons and the recurrent E/I assemblies cause a temporal delay of Type 2 inhibition with respect to feed-forward input currents. To compensate for this delay in the continuous-time version BCP, we implemented short-term filtering of feed-forward synaptic currents and presynaptic activity. The resulting learning rule is:

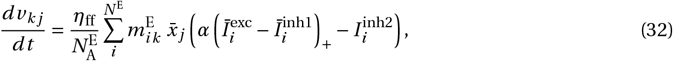

where short-term moving averages of the feed-forward currents were implemented as leaky integrators

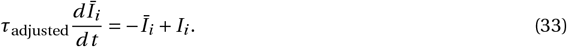

Here we choose the filtering time constant simply as

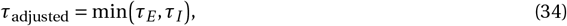

which matches the fastest network mode under slow inputs (see Supplementary Material 1.4).

Similarly, the presynaptic activity is filtered to obtain the moving average 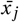 using a leaky integrator with time constant *τ*_pre_:

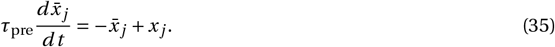

This temporal filtering of both synaptic currents and presynaptic inputs reduces the temporal mismatch between Type 2 inhibitory currents and feed-forward inputs, thereby stabilizing online learning.

#### Plasticity at output neurons

We parameterize the readout weights from excitatory neuron *j* to readout neuron *i* as

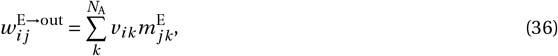

where *v*_*ik*_ is the element *i, k* of the readout weight matrix 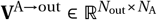. To update readout weights during learning of target **y**, we used a simple Delta rule [65] multiplying the presynaptic activity by the difference between the target output and the actual output:

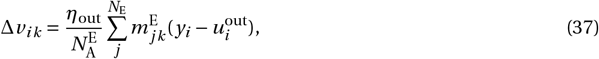

where *η*_out_ is the learning rate for the readout weights. This learning rule is equivalent to the gradient of the squared error loss function 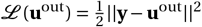 with respect to the readout weights.

### Simulation details

All numerical simulations were implemented using custom software written in Python 3.11.8 and executed on NVIDIA Quadro RTX 5000 GPUs. Software packages employed for numerical simulations included Jax [137], Flax [138], and Diffrax [139]. The Fashion-MNIST dataset was obtained from the TensorFlow datasets library [140]. The code to reproduce our results is available at https://github.com/fmi-basel/balance-controlled-plasticity. All hyperparameters for numerical simulations can be found in Supplementary Tables S1-S5.

#### Single-assembly experiments

To investigate the learning dynamics in a single assembly (Figs. 2 and 3), we simulated an assembly comprising 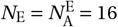 excitatory neurons and 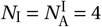 inhibitory neurons.

#### Dynamic input stimuli

Assembly neurons received feed-forward input from *N*_X_ *=* 20 input neurons, each encoding a linear combination of *N*_S_ *=* 3 dynamic stimuli (A, B, and C). The temporal activity of each stimulus was modeled as a rectified Ornstein-Uhlenbeck (OU) process. Specifically, for each stimulus *i*, we simulated a standard OU process *z*_*i*_ (*t*) according to

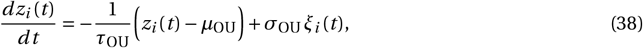

where ξ_*i*_ (*t*) is Gaussian white noise with zero mean and unit variance, *τ*_OU_ is the OU time constant, *µ*_OU_ is the process mean, and *σ*_OU_ is the fluctuation magnitude. The effective stimulus activity was obtained by thresholding:

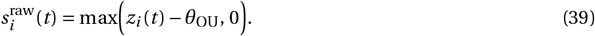

To generate smooth input signals, the thresholded activity was low-pass filtered using an exponential filter with time constant *τ*_filter_, resulting in the corresponding stimulus dynamics:

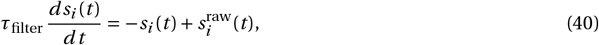

with initial condition 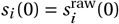.

#### Stimulus encoding

Input neurons encoded the stimuli by forming linear combinations based on a Gaussian tiling of the stimulus space with circular boundaries. The activity of input neuron *x*_*i*_ (*t*) was given by

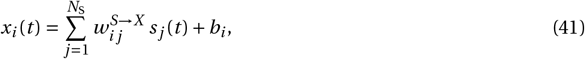

where 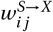 denotes the synaptic weight from stimulus *j* to input neuron *i*, and *b*_*i*_ is a bias term drawn from a standard lognormal distribution scaled by *b*_scale_. The synaptic weights **W**^*S→X*^ were determined using a Gaussian tiling scheme. In particular, the weight from stimulus *j* to input neuron *i* was given by

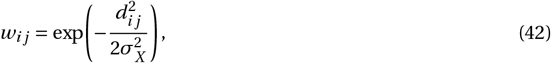

where *σ*_*X*_ is the tuning width parameter and *d*_*i j*_ denotes the circular distance between the preferred tuning Λ_*i*_ of input neuron *i* and the stimulus location *ξ*_*j*_. The circular distance was defined as

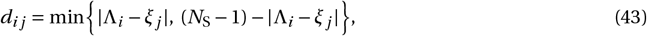

with preferred tunings

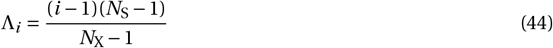

and stimulus locations *ξ*_*j*_ *= j −* 1. This construction guarantees that the preferred tunings of input neurons are distributed uniformly over a circular domain.

#### Weight initialization

The feed-forward weights **V**^*X →A*^ were initialized from a normal distribution with mean 0.2 and standard deviation 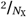. The output weights were set as

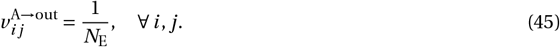

#### Numerical integration

Input stimulus activity was generated in 15-s chunks using the forward Euler method with a step size of Δ*t =* 1 ms. Linear interpolation between time steps was used to produce continuous signals. The network dynamics were then integrated using Tsitouras’ 5th order explicit Runge-Kutta method [141] with an adaptive step size. Simulations were run continuously for the duration of each experiment.

#### Manual feedback perturbations

For the manual feedback perturbation experiments (Fig. 2d), network dynamics were simulated for *T =* 15 s while applying two sets of strong external current perturbations *I* ^ext^ to all inhibitory neurons in the assembly. The dynamics of the inhibitory membrane potential were governed by

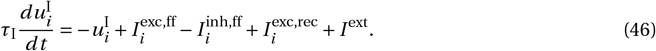

Two perturbation phases were administered, each lasting 3 s and separated by a 3-s interval. In the first phase, a positive perturbation of *I* ^ext^ *=* 10 was applied, while in the second phase a negative perturbation of *I* ^ext^ *= −*10 was applied.

#### Trajectory matching task

For the trajectory matching task (see Figs. 4 and 5), we simulated networks with *N*_*X*_ *=* 20 input neurons, a hidden layer consisting of *N*_*E*_ *=* 240 excitatory and *N*_*I*_ *=* 60 inhibitory neurons arranged in *N*_*A*_ *=* 20 assemblies, and an output layer with *N*_Y_ *=* 2 output neurons.

#### Periodic inputs

Input neurons encoded random linear combinations of 20 sinusoidal functions. The activity of the *j* th sinusoidal function was defined as

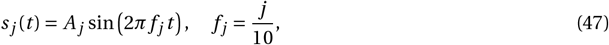

where *A*_*j*_ is a random amplitude drawn from a uniform distribution between 0.1 and 1.0 and *f* _*j*_ is the frequency of the sinusoidal function, parameterized so that each sinusoidal function completes an integer number of periods every 10 seconds. Each input neuron encoded a linear combination of sinusoidal functions so that the activity of the *i* th input neuron was given by

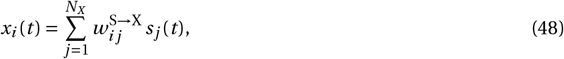

where 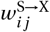 are the elements of a mixing matrix **W**^S*→*X^ drawn from a standard normal distribution.

#### Target trajectory

The target trajectory was generated from a hand-drawn turtle-shaped vector path in the 2D plane. We obtained the target **y** by sampling 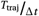 uniformly spaced points along the path, where Δ*t* is the simulation time step and *T*_traj_ *=* 5 seconds is the total duration of the trajectory. The trajectory was centered at the origin and scaled to lie within [*−*1, 1] in both the *X* and *Y* dimensions.

#### Parameter initialization

Feed-forward weights **V**^X*→*A^ were initialized from a normal distribution with mean 0 and standard deviation 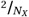. Similarly, readout weights **V**^A*→*out^ were initialized from a normal distribution with mean 0 and standard deviation 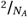.

#### Training

Networks were trained to minimize the mean squared error (MSE) between the network output and the target trajectory. Top-down feedback was applied via a proportional controller with fixed gain *k*_*p*_ through feedback weights **Q** defined in equation (29). The network was trained for a total duration of *T =* 360 minutes, corresponding to 4320 iterations of the trajectory. Synaptic weights were recorded every 20 iterations during training. Numerical integration was performed using the forward Euler method with a time step of Δ*t =* 1 ms. To prevent runaway LTD due to excess inhibition in the absence of postsynaptic activity in networks with assembly overlap (Fig. 5), we constrained all synaptic weights to the interval [*−*1, 1] during training.

#### Assessing model performance

Model performance was evaluated under two conditions: with top-down feedback (closed-loop) and without top-down feedback (open-loop). Closed-loop performance was measured from the recorded network output every 20 iterations. Since open-loop performance cannot be directly observed when top-down feedback is active, we created a copy of the model every 20 iterations, disabled top-down feedback, and recorded the open-loop output for 2 iterations. Performance was quantified using the MSE and the coefficient of determination (*R*^2^) between the network output and the target trajectory.

#### Control experiments

Two control conditions were examined: one without top-down feedback and one without a hidden layer. In the no-feedback condition, feedback weights **Q** were set to zero, effectively disabling top-down feedback while still optimizing the readout weights to minimize the MSE. In the no-hidden layer condition, the hidden layer was removed and the input neurons were connected directly to the output neurons, with the readout weights trained normally.

### Training deep hierarchical networks on Fashion-MNIST

We rescaled all Fashion-MNIST input images to values between 0 and 1 and flattened each image into a 784-dimensional vector that served as the network input. Each hidden layer consisted of *N*_A_ *=* 128 E/I assemblies, with each assembly containing 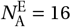 excitatory and 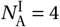 inhibitory neurons. The output layer comprised *N*_Y_ *=* 10 neurons, one for each class in the Fashion-MNIST dataset. For networks with multiple hidden layers, each hidden layer was treated as a separate E/I network with dynamics defined in equations (14) and (15). The first hidden layer received the image input *x*, and subsequent layers were connected sequentially, with the output of one layer serving as the input to the next. The final hidden layer was connected to the output layer.

#### Top-down feedback

Each hidden layer *l ∈* {1, 2,…, *L*} received direct top-down feedback from a controller. Feedback weights followed the mapping defined in equation (29) adjusted for multiple hidden layers, so that the feedback weights to the *l* th hidden layer were given by

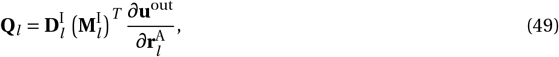

where 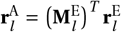 represents the activity of each assembly in the *l* th hidden layer. For networks with three hidden layers, we additionally implemented a layer-wise normalization procedure by dividing each layer’s feedback weight matrix **Q**_*l*_ by its Frobenius norm ∥**Q**_*l*_ ∥_*F*_. This normalization, while not required for successful training, significantly accelerated the network’s convergence to equilibrium, and reduced variance in performance across training runs, resulting in more reliable network behavior.

#### Classification loss function

For the classification task, we adapted the standard cross-entropy loss function. Conventionally, the output layer employs a Softmax activation to match a one-hot encoded target vector **y**. However, since softmax outputs can cause issues in feedback control settings [81], we used soft targets instead of one-hot encodings: the target value was set to 0.99 for the correct class and to 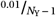 for all other classes. In line with previous work [81, 125], we utilized a linear readout layer and incorporated the softmax operation within the loss function:

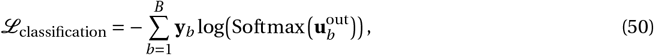

where *B* is the batch size and 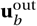 denotes the network output for the *b*th sample.

#### Parameter initialization

Network parameters were initialized from normal distributions with mean 0 and standard deviation 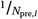, where *N*_pre,*l*_ is the number of neurons in the preceding layer.

#### Training

Networks were trained for 50 epochs using a batch size of *B =* 100. For each sample, the network was first allowed to converge to an open-loop equilibrium state in the absence of top-down feedback.

Numerical integration was performed using Tsitouras’ 5th order explicit Runge-Kutta method [141] with an adaptive step size, running for up to 2 seconds or until an equilibrium state was reached (defined by an absolute tolerance of 10^*−*6^). At the open-loop equilibrium state, feedback weights **Q** were computed.

Thereafter, top-down feedback control was enabled, and the network was allowed to converge to a closed-loop equilibrium state using the same integration method. Parameter updates were then computed at the closed-loop equilibrium state using the BCP learning rule in equation (31). We constrained all parameters to the interval [*−*1, 1] during training to prevent extreme values in the network Jacobian that could arise from excessively large weights. All simulations employed the ADAM optimizer with a learning rate of 0.001.

#### Backpropagation control experiments

To compare the performance of the BCP learning rule with standard backpropagation, we trained networks of identical architecture using standard backpropagation. To match the number of parameters in the E/I assembly networks, the number of hidden units was set to *N =* 128. As in the BCP networks, all hidden units used a rectified linear activation function. We trained networks with either 1 or 3 hidden layers using the ADAM optimizer (learning rate 0.001, batch size 100) for50 epochs. In contrast to the BCP networks, these networks employed a softmax output layer and a standard cross-entropy loss function.

### Model of learning in BLA during fear conditioning

To model the fear conditioning experiments by Krabbe et al. [46], we simulated networks with *N*_X_ *=* 20 input neurons, a hidden layer consisting of *N*_E_ *=* 160 excitatory and *N*_I_ *=* 40 inhibitory neurons arranged in *N*_A_ *=* 40 assemblies, and a single readout neuron that was used to model freezing probability in response to the CS. Networks were trained on a total of ten fear conditioning trials, each trial lasting 20 seconds. During each trial, the CS was presented for a duration of 4 seconds, followed by a US presentation for 2 seconds and an inter-trial interval period of 14 seconds.

#### CS encoding and inputs

The activity of input neurons was modeled as *x*_*j*_ (*t*) *= ϵ*_*j*_ (*t*) *+r*_*j*_ (*t*), where *ϵ*_*j*_ (*t*) is random noise determined by an OU process with parameters *τ*_OU_, *µ*_OU_, and *σ*_OU_, and *r* _*j*_ (*t*) represents the filtered response of neuron *j* to the CS. The response selectivity of inputs to the CS was chosen to generate a sparse encoding. Specifically, we randomly selected 25% of the input neurons to respond to the CS with response strengths *q*_*j*_ drawn from a lognormal distribution and normalized to sum to 1. The raw binary CS presentations were filtered with an alpha kernel with separate rise (*τ*_rise_) and decay (*τ*_decay_) time constants to produce temporally smoothed responses *r* _*j*_ (*t*), calculated as:

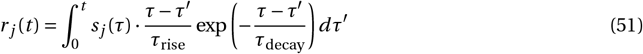

where *s* _*j*_ (*t*) is a binary input signal for neuron *j* that represents CS presentation.

#### Freezing probability

We used a single readout neuron representing a continuous valence prediction output of the BLA. Following experimental evidence for genetically identifiable subpopulations of BLA neurons for positive and negative valence encoding [142], we used random, fixed readout weights, so that a negative (positive) weight indicated an assembly encoding negative (positive) valence. Freezing probability was modeled as a shifted logistic function operating on the valence prediction:

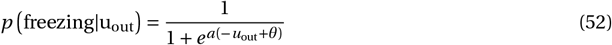

#### US-driven feedback and loss function

We modeled the US as an external stimulus that sets a target freezing probability of 1 for the network. During US presentation, a top-down controller as described above provided feedback to all hidden layer Type 2 interneurons. Top-down feedback was set to zero in the absence of the US.

### Model of motor learning in M1

To model the motor learning experiments by Ren et al. [47], we trained networks to output a motor command representing a lever push-and-pull movement in response to a sensory cue. We simulated networks with *N*_*X*_ *=* 20 input neurons, a hidden layer consisting of *N*_*E*_ *=* 320 excitatory and *N*_*I*_ *=* 80 inhibitory neurons arranged in *N*_*A*_ *=* 80 assemblies, and a single readout neuron representing a motor command. Networks were trained for a total of 2000 trials, each lasting 8 seconds. Two seconds into each trial, the cue was presented and the network was tasked to generate the appropriate motor command.

#### Cue encoding and inputs

The cue signal was designed to capture the transient non-normal amplification dynamics observed in motor cortex [143]. Specifically, we modeled the cue-evoked neuronal response as an alpha kernel with rise and decay time constants *τ*_rise_ and *τ*_decay_ that provided a transient temporal envelope. Each neuron’s response to the cue within this temporal window was modeled as a linear combination of *N*_sines_ sinusoidal waves with frequencies randomly sampled between 0.1 and 2.0 Hz and random phases. This was implemented as:

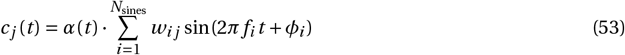

where *w*_*i j*_ are elements of a random weight matrix drawn from a normal distribution and scaled by a factor of 10, and *α*(*t*) is the normalized alpha kernel. The resulting cue activity began 2 seconds into each trial and was normalized to have a maximum amplitude of 1. Input neuron activity was modeled as *x*_*j*_ (*t*) *= c* _*j*_ (*t*)*+ϵ*_*j*_ (*t*), where *c* _*j*_ (*t*) is the filtered cue input to neuron *j* and *ϵ*_*j*_ (*t*) is random noise determined by an OU process with parameters *τ*_OU_, *µ*_OU_, and *σ*_OU_.

#### Motor output

The target motor command was defined as a single sinusoidal cycle that began at time *t*_cue_ *=* 2 seconds and lasted for a duration of *T*_movement_ *=* 2 seconds:

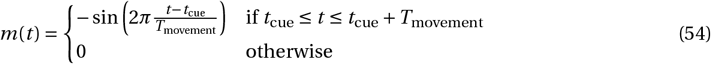

The network output was trained to follow the motor command *m*(*t*) given the cue inputs like described previously for the trajectory matching task.

#### Lever output and correct trials

The lever position was modeled as the temporal integration of the motor command, with an offset correction to ensure the lever position started at zero at cue onset:

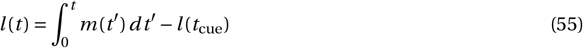

Following Ren et al. [47], we classified trials as successful when the lever trajectory crossed a threshold *θ*_lever_ both downward and upward within 2 seconds after cue onset, corresponding to a complete push-pull movement.

