## Supplementary Material for "Breaking Balance: Encoding local error signals in perturbations of excitation-inhibition balance"

### 1 Supplementary Methods

#### 1.1 Optimization of Type 2 inhibitory synaptic connections for precise E/I balance

Throughout this paper, we require that the Type 2 (recurrent) inhibitory currents maintain a proportional relationship with the total feed-forward input. For each excitatory neuron $i$, this balance condition is expressed as (c.f. Eq. (2)):

$$I_i^{\mathrm{inh2}}(t) = \alpha\left(\bar{I}_i^{\mathrm{exc}}(t) - \bar{I}_i^{\mathrm{inh1}}(t)\right)_+ \tag{S1}$$

where $\alpha < 1.0$ is a proportionality parameter.

We directly optimized the I → E synaptic weights to satisfy this balance condition. To this end, we assume constant inputs to the network $\mathbf{x}$, which allows us to approximate neuronal responses by their steady-state values. Setting the time derivatives in equations (14) and (15) to zero and using the feed-forward weight parameterization from equation (19), we obtain:

$$\begin{aligned} \overset{*}{\mathbf{u}}^{\mathrm{E}} &= \mathbf{M}^{\mathrm{E}}\mathbf{V}^{\mathrm{X}\to\mathrm{A}}\mathbf{x} - \mathbf{W}^{\mathrm{I}\to\mathrm{E}}\overset{*}{\mathbf{r}}^{\mathrm{I}}, & \text{(S2)}\\ \overset{*}{\mathbf{u}}^{\mathrm{I}} &= \mathbf{M}^{\mathrm{I}}\mathbf{V}^{\mathrm{X}\to\mathrm{A}}\mathbf{x} + \mathbf{W}^{\mathrm{E}\to\mathrm{I}}\overset{*}{\mathbf{r}}^{\mathrm{E}}. & \text{(S3)}\end{aligned}$$

**Linearized Dynamics and Loss Function.** Since precise E/I balance is only relevant when neurons are active, we restrict the optimization to the active regime[1]. Formally, let

$$\mathscr{I} = \{i : \left[\mathbf{M}^{\mathrm{E}}\mathbf{V}^{\mathrm{X}\to\mathrm{A}}\mathbf{x}\right]_i > 0\} \tag{S4}$$

denote the set of indices corresponding to neurons with strictly positive feed–forward input. Because we use a rectified-linear activation $r_i = \mathrm{ReLU}(u_i) = \max(0, u_i)$, for any neuron $i$ with strictly positive feed-forward drive,

$$[\mathbf{M}^{\mathrm{E}}\mathbf{V}^{\mathrm{X}\to\mathrm{A}}\mathbf{x}]_i > 0 \quad\Longrightarrow\quad r_i = u_i. \tag{S5}$$

Hence, we replace $\overset{*}{\mathbf{r}}^E$ and $\overset{*}{\mathbf{r}}^I$ by $\overset{*}{\mathbf{u}}^E$ and $\overset{*}{\mathbf{u}}^I$, respectively, to reduce the steady-state dynamics to a linear system:

$$\begin{aligned} \overset{*}{\mathbf{u}}^{\mathrm{E}} &= \mathbf{M}^{\mathrm{E}}\mathbf{V}^{\mathrm{X}\to\mathrm{A}}\mathbf{x} - \mathbf{W}^{\mathrm{I}\to\mathrm{E}}\overset{*}{\mathbf{u}}^{\mathrm{I}}, & \text{(S6)}\\ \overset{*}{\mathbf{u}}^{\mathrm{I}} &= \mathbf{M}^{\mathrm{I}}\mathbf{V}^{\mathrm{X}\to\mathrm{A}}\mathbf{x} + \mathbf{W}^{\mathrm{E}\to\mathrm{I}}\overset{*}{\mathbf{u}}^{\mathrm{E}}. & \text{(S7)}\end{aligned}$$

Substituting the first equation into the second yields

$$\begin{aligned} \overset{*}{\mathbf{u}}^{\mathrm{I}} &= \mathbf{M}^{\mathrm{I}}\mathbf{V}^{\mathrm{X}\to\mathrm{A}}\mathbf{x} + \mathbf{W}^{\mathrm{E}\to\mathrm{I}}\left(\mathbf{M}^{\mathrm{E}}\mathbf{V}^{\mathrm{X}\to\mathrm{A}}\mathbf{x} - \mathbf{W}^{\mathrm{I}\to\mathrm{E}}\overset{*}{\mathbf{u}}^{\mathrm{I}}\right)\\ &= \left(\mathbf{M}^{\mathrm{I}} + \mathbf{W}^{\mathrm{E}\to\mathrm{I}}\mathbf{M}^{\mathrm{E}}\right)\mathbf{V}^{\mathrm{X}\to\mathrm{A}}\mathbf{x} - \mathbf{W}^{\mathrm{E}\to\mathrm{I}}\mathbf{W}^{\mathrm{I}\to\mathrm{E}}\overset{*}{\mathbf{u}}^{\mathrm{I}}. & \text{(S8)}\end{aligned}$$

Rearranging, we obtain

$$\left(I + \mathbf{W}^{\mathrm{E}\to\mathrm{I}}\mathbf{W}^{\mathrm{I}\to\mathrm{E}}\right)\overset{*}{\mathbf{u}}^{\mathrm{I}} = \left(\mathbf{M}^{\mathrm{I}} + \mathbf{W}^{\mathrm{E}\to\mathrm{I}}\mathbf{M}^{\mathrm{E}}\right)\mathbf{V}^{\mathrm{X}\to\mathrm{A}}\mathbf{x}, \tag{S9}$$

so that

$$\overset{*}{\mathbf{u}}^{\mathrm{I}} = \left(I + \mathbf{W}^{\mathrm{E}\to\mathrm{I}}\mathbf{W}^{\mathrm{I}\to\mathrm{E}}\right)^{-1}\left(\mathbf{M}^{\mathrm{I}} + \mathbf{W}^{\mathrm{E}\to\mathrm{I}}\mathbf{M}^{\mathrm{E}}\right)\mathbf{V}^{\mathrm{X}\to\mathrm{A}}\mathbf{x}. \tag{S10}$$

We now link this expression to the balance condition in Equation (2). In vectorized form, the balance condition can be formulated as

$$I_i^{\mathrm{inh2}} = \alpha\left[\mathbf{M}^{\mathrm{E}}\mathbf{V}^{\mathrm{X}\to\mathrm{A}}\mathbf{x}\right]_i. \tag{S11}$$

Noting that the Type 2 inhibitory current is given by $\mathbf{W}^{\mathrm{I}\to\mathrm{E}}\overset{*}{\mathbf{u}}^{\mathrm{I}}$, we combine equations (S10) and (S11) to yield

$$\mathbf{W}^{\mathrm{I}\to\mathrm{E}}\left(I + \mathbf{W}^{\mathrm{E}\to\mathrm{I}}\mathbf{W}^{\mathrm{I}\to\mathrm{E}}\right)^{-1}\left(\mathbf{M}^{\mathrm{I}} + \mathbf{W}^{\mathrm{E}\to\mathrm{I}}\mathbf{M}^{\mathrm{E}}\right)\mathbf{V}^{\mathrm{X}\to\mathrm{A}}\mathbf{x} = \alpha\mathbf{M}^{\mathrm{E}}\mathbf{V}^{\mathrm{X}\to\mathrm{A}}\mathbf{x}, \tag{S12}$$

[1] In non-overlapping assemblies, if the feed–forward inputs to the assembly are negative, then both excitatory and inhibitory neurons are silent and the EI balance condition is trivially satisfied because $I_i^{\mathrm{inh2}} = 0$ for all neurons in that assembly. Note that, for overlapping circuits, this assumption is not strictly valid and introduces a potential systematic issue because neurons can receive inhibitory currents unrelated to the feed–forward input from that assembly, leading to systematic deviations from the desired EI balance target (as illustrated in Fig. 5).

which must hold for all active neurons. Since $\mathbf{V}^{X\to A}\mathbf{x}$ appears on both sides, it can be factored out, yielding the parameter condition

$$\mathbf{W}^{I\to E}\left(I+\mathbf{W}^{E\to I}\mathbf{W}^{I\to E}\right)^{-1}\left(\mathbf{M}^{I}+\mathbf{W}^{E\to I}\mathbf{M}^{E}\right)=\alpha\mathbf{M}^{E}. \tag{S13}$$

Defining the residual matrix $\mathbf{R}$ as the difference between the two sides of (S13),

$$\mathbf{R}=\mathbf{W}^{I\to E}\left(I+\mathbf{W}^{E\to I}\mathbf{W}^{I\to E}\right)^{-1}\left(\mathbf{M}^{I}+\mathbf{W}^{E\to I}\mathbf{M}^{E}\right)-\alpha\mathbf{M}^{E}, \tag{S14}$$

we cast our goal of precise E/I balance as the following optimization problem over the active regime:

$$\min_{\mathbf{W}^{I\to E}}\mathscr{L}_{\text{balance}}\left(\mathbf{W}^{I\to E}\right)\quad\text{with}\quad\mathscr{L}_{\text{balance}}\left(\mathbf{W}^{I\to E}\right)=\frac{1}{n}\|\mathbf{R}\left(\mathbf{W}^{I\to E}\right)\|_F^2, \tag{S15}$$

where $n$ denotes the total number of entries in $\mathbf{R}$ and $\|\cdot\|_F$ denotes the Frobenius norm.

**Optimization Procedure.** To optimize $\mathbf{W}^{I\to E}$, we first mapped an unconstrained parameter matrix (denoted by $\mathbf{P}$) to strictly positive inhibitory-to-excitatory synaptic weights via

$$\mathbf{W}^{I\to E}=\text{ReLU}(\mathbf{P}). \tag{S16}$$

Assuming that the $\mathbf{W}^{I\to E}$ weights are approximately proportional to the corresponding $\mathbf{W}^{E\to I}$ weights, we initialize

$$\mathbf{P}_{\text{init}}=\alpha\left(\mathbf{W}^{E\to I}\right)^{T}, \tag{S17}$$

so that the initial inhibitory current roughly matches $\alpha\mathbf{M}^{E}\mathbf{V}\mathbf{x}$. We then minimize the loss (S15) with respect to $\mathbf{P}$ using RMSprop with a cosine decay schedule. In each optimization step, we compute the gradient

$$\nabla_{\mathbf{P}}\mathscr{L}\left(\mathbf{W}^{I\to E}(\mathbf{P})\right), \tag{S18}$$

update $\mathbf{P}$, and enforce nonnegativity via the ReLU mapping. For numerical stability, we added a small regularization term $\delta=10^{-6}$ to the identity in Equation (S14) when computing the inverse. For each network, we performed the optimization for 20000 steps with a learning rate of $10^{-3}$, and set the final inhibitory weight matrix to the optimized $\mathbf{W}^{I\to E}$.

#### 1.2 Feedback weights approximate network Jacobian.

Learning algorithms based on minimization of top-down control inputs rely on top-down feedback that can minimize a network output error. Meulemans et al. [9, 81, 87] have shown that a linear controller $\mathbf{W}^{I\to E}\mathbf{Q}\mathbf{c}$ can be sufficient to minimize the loss function $\mathscr{L}(\mathbf{u}^{\text{out}})$, provided that the column space of the feedback weights is equal to the row space of the network Jacobian $\mathbf{J}$ at steady-state [9, 81]. The network Jacobian for networks with a single hidden layer in our rate-based framework can be formally defined as

$$\mathbf{J}=\frac{\partial\mathbf{u}^{\text{out}}}{\partial\mathbf{u}^{E}} \tag{S19}$$

and describes the effect of small perturbations in the excitatory membrane potentials on the network output $\mathbf{u}^{\text{out}}$.

The feedback mapping defined in (29) is not exactly equal to the transpose of the network Jacobian. However, it is easy to show that the column space condition is fulfilled for this choice of feedback weights. Specifically, we require that

$$\text{col}\left(\mathbf{W}^{I\to E}\mathbf{Q}\right)\quad=\quad\text{col}\left(\frac{\partial\mathbf{u}^{\text{out}}}{\partial\mathbf{u}^{E}}\right)^{T}. \tag{S20}$$

Substituting the feedback weights (29) into the left-hand side and evaluating the derivative on the right, we need to show that

$$\text{col}\left(\mathbf{W}^{I\to E}\mathbf{D}^{I}\left(\mathbf{M}^{I}\right)^{T}\left(\mathbf{V}^{A\to\text{out}}\right)^{T}\right)=\text{col}\left(\mathbf{D}^{E}\left(\mathbf{M}^{E}\right)^{T}\left(\mathbf{V}^{A\to\text{out}}\right)^{T}\right), \tag{S21}$$

where $\mathbf{D}^{E}$ is a diagonal matrix containing the derivative of the rectified nonlinearity for each neuron, i.e. $\mathbf{D}^{E}_{ii}=\phi'(u^{E}_{i})$. The inhibitory-to-excitatory weights $\mathbf{W}^{I\to E}$ are optimized to yield proportional balance, which is achieved through reciprocal connectivity with other excitatory neurons in the same assembly. It is thus reasonable to assume that column space of $\mathbf{W}^{\mathrm{I}\to\mathrm{E}}$ is equal to the row space of the reciprocal excitatory-to-inhibitory weights $\left(\mathbf{W}^{\mathrm{E}\to\mathrm{I}}\right)^T$. This is always true at initialization and for non-overlapping assembly structures. While we cannot guarantee that this condition holds for arbitrary assembly structures, we find this to be true empirically in our numerical simulations. Thus, assuming $\mathrm{col}\left(\mathbf{W}^{\mathrm{I}\to\mathrm{E}}\right) = \mathrm{col}\left(\left(\mathbf{W}^{\mathrm{E}\to\mathrm{I}}\right)^T\right)$ and given the definition in of $\mathbf{W}^{\mathrm{E}\to\mathrm{I}}$ in (17), we have

$$\mathrm{col}\left(\mathbf{W}^{\mathrm{I}\to\mathrm{E}}\right) = \mathrm{col}\left(\left(\mathbf{M}^{\mathrm{E}}\right)^T \mathbf{M}^{\mathrm{I}}\right). \tag{S22}$$

Moreover, since the assembly structure enforces that neurons within the same assembly share identical feed-forward tuning, we can reasonably assume that the derivative terms projected into assembly space, $\mathbf{M}^{\mathrm{E}}\mathbf{D}^{\mathrm{E}}$ and $\mathbf{M}^{\mathrm{I}}\mathbf{D}^{\mathrm{I}}$, span the same subspace, i.e.

$$\mathrm{col}\left(\mathbf{M}^{\mathrm{E}}\mathbf{D}^{\mathrm{E}}\right) = \mathrm{col}\left(\mathbf{M}^{\mathrm{I}}\mathbf{D}^{\mathrm{I}}\right). \tag{S23}$$

Consequently, there exists an invertible transformation $\mathbf{C}$ such that

$$\mathbf{M}^{\mathrm{I}}\mathbf{D}^{\mathrm{I}} = \mathbf{M}^{\mathrm{E}}\mathbf{D}^{\mathrm{E}}\mathbf{C}. \tag{S24}$$

Substituting these relationship into (S21), we find

$$\begin{aligned}
\mathrm{col}\left(\mathbf{W}^{\mathrm{I}\to\mathrm{E}}\mathbf{D}^{\mathrm{I}}\left(\mathbf{M}^{\mathrm{I}}\right)^T\left(\mathbf{V}^{\mathrm{A}\to\mathrm{out}}\right)^T\right) &= \mathrm{col}\left(\left(\mathbf{M}^{\mathrm{E}}\right)^T\mathbf{M}^{\mathrm{I}}\mathbf{D}^{\mathrm{I}}\left(\mathbf{M}^{\mathrm{I}}\right)^T\left(\mathbf{V}^{\mathrm{A}\to\mathrm{out}}\right)^T\right) && \text{(S25)}\\
&= \mathrm{col}\left(\left(\mathbf{M}^{\mathrm{E}}\right)^T\mathbf{M}^{\mathrm{E}}\mathbf{D}^{\mathrm{E}}\mathbf{C}\left(\mathbf{M}^{\mathrm{I}}\right)^T\left(\mathbf{V}^{\mathrm{A}\to\mathrm{out}}\right)^T\right) && \text{(S26)}\\
&= \mathrm{col}\left(\mathbf{D}^{\mathrm{E}}\left(\mathbf{M}^{\mathrm{E}}\right)^T\left(\mathbf{V}^{\mathrm{A}\to\mathrm{out}}\right)^T\right), && \text{(S27)}
\end{aligned}$$

which completes the proof.

##### 1.3 Derivation of the BCP learning rule

The BCP learning framework stipulates that weight changes are driven by deviations from the proportional E/I balance target described in Equation (2). Here, we show how this learning rule can be derived from first principles for the rate-based framework described above.

**Learning as Minimization of Top-Down Feedback Control.** Recent theoretical frameworks have established a *least control* principle as an effective learning strategy for artificial neural networks that operate at an equilibrium state [81, 87]. These frameworks assume an arbitrary system with state $\mathbf{u}$ and parameters $\theta$, with dynamics given by

$$\tau\frac{d\mathbf{u}}{dt} = f(\mathbf{u},\theta) + \psi(t), \tag{S28}$$

where $\psi(t)$ is a control signal that drives the system to a desired steady-state $\overset{*}{\mathbf{u}}$ at which a loss function $\mathcal{L}(\overset{*}{\mathbf{u}})$ is minimized, i.e. $f(\overset{*}{\mathbf{u}},\theta) = 0$ and $\nabla_{\mathbf{u}}\mathcal{L}(\overset{*}{\mathbf{u}}) = 0$. In this system, learning can be re-formulated as a minimization of control problem [87]: Introducing a surrogate loss $\mathcal{H}(\theta) = ||\psi||^2$ for the control signal, the learning rule can be derived as a gradient-following update in $\theta$:

$$\nabla_\theta\mathcal{H}(\theta) = -\left(\frac{\partial f(\mathbf{u},\theta)}{\partial\theta}\right)^T \overset{*}{\psi}. \tag{S29}$$

Note that this framework relies on two key assumptions: (1) the control signal $\psi$ exerts a linear influence on the system state, and (2) the control is optimal in the sense that it minimizes the loss $\mathcal{L}(\overset{*}{\mathbf{u}})$.

**Derivation of BCP as minimizing top-down feedback control.** We start from the vectorized dynamics of the rate-based E/I network and make the linear top-down feedback input explicit (c.f. Eqs. (14) and (15)):

$$\begin{aligned}
\tau_{\mathrm{E}}\frac{d\mathbf{u}^{\mathrm{E}}}{dt} &= -\mathbf{u}^{\mathrm{E}} + \mathbf{W}^{\mathrm{X}\to\mathrm{E}}\mathbf{x} - \mathbf{W}^{\mathrm{I}\to\mathrm{E}}\mathbf{r}^{\mathrm{I}}, && \text{(S30)}\\
\tau_{\mathrm{I}}\frac{d\mathbf{u}^{\mathrm{I}}}{dt} &= -\mathbf{u}^{\mathrm{I}} + \mathbf{W}^{\mathrm{X}\to\mathrm{I}}\mathbf{x} + \mathbf{W}^{\mathrm{E}\to\mathrm{I}}\mathbf{r}^{\mathrm{E}} + \mathbf{Q}\,\mathbf{c}, && \text{(S31)}
\end{aligned}$$

where $\mathbf{Q}$ is the matrix of top-down feedback weights and $\mathbf{c}$ denotes the activity of a neuronal population exerting feedback control. Under the assumption that inhibitory dynamics are much faster than excitatory ones ($\tau_I \ll \tau_E$), we set

$$\tau_I \frac{d\mathbf{u}^I}{dt} \approx 0, \tag{S32}$$

which yields a quasi–steady state for the inhibitory firing rates:

$$\mathbf{r}^I = \phi\Big(\mathbf{W}^{X\to I}\mathbf{x} + \mathbf{W}^{E\to I}\phi(\mathbf{u}^E) + \mathbf{Q}\mathbf{c}\Big). \tag{S33}$$

Substituting this expression back into the excitatory dynamics, we express the dynamics of the network as a single ODE for the excitatory membrane potentials:

$$\tau_E \frac{d\mathbf{u}^E}{dt} = -\mathbf{u}^E + \mathbf{W}^{X\to E}\mathbf{x} - \mathbf{W}^{I\to E}\phi\Big(\mathbf{W}^{X\to I}\mathbf{x} + \mathbf{W}^{E\to I}\phi(\mathbf{u}^E) + \mathbf{Q}\mathbf{c}\Big). \tag{S34}$$

In networks operating in the precisely balanced regime defined in (S11), the inhibitory currents are proportional to the feed-forward excitatory input. Specifically, in the absence of top-down feedback (i.e. when $\mathbf{Q}\mathbf{c} = \mathbf{0}$), precise balance implies that:

$$\mathbf{W}^{I\to E}\mathbf{r}^I = \begin{cases} \alpha\mathbf{W}^{X\to E}\mathbf{x}, & \text{if } \mathbf{W}^{X\to E}\mathbf{x} > \mathbf{0}, \\ \mathbf{0}, & \text{if } \mathbf{W}^{X\to E}\mathbf{x} \le \mathbf{0}, \end{cases} \tag{S35}$$

so that, in the active regime with positive feed-forward inputs, inhibitory currents are proportional to the balance target. Thus, assuming assuming the network is in the active regime and the EI balance condition is fulfilled, the dynamics of our network in the absence of top-down feedback can be simplified to

$$\tau_E \frac{d\mathbf{u}^E}{dt} = -\mathbf{u}^E + \mathbf{W}^{X\to E}\mathbf{x} - \alpha\mathbf{W}^{X\to E}\mathbf{x}, \qquad \mathbf{Qc} = \mathbf{0}. \tag{S36}$$

We now consider the effect of top-down feedback and assume that the top-down feedback only influences inhibitory interneurons that are active, i.e. $\mathbf{Qc} = \mathbf{0}$ if $\mathbf{r}^I = \mathbf{0}$. Because those interneurons that are active are operating in a linear regime, we can reasonably assume a linear effect of the feedback on the inhibitory activity, so that

$$\mathbf{W}^{I\to E}\mathbf{r}^I = \alpha\mathbf{W}^{X\to E}\mathbf{x} + \mathbf{W}^{I\to E}\mathbf{Qc}. \tag{S37}$$

and approximate the dynamics of the network as

$$\begin{aligned} \tau_E \frac{d\mathbf{u}^E}{dt} &= -\mathbf{u}^E + \mathbf{W}^{X\to E}\mathbf{x} - \alpha\mathbf{W}^{X\to E}\mathbf{x} + \mathbf{W}^{I\to E}\mathbf{Qc}, & \text{(S38)} \\ &= -\mathbf{u}^E + (1-\alpha)\mathbf{W}^{X\to E}\mathbf{x} + \mathbf{W}^{I\to E}\mathbf{Qc}. & \text{(S39)} \end{aligned}$$

which can be naturally mapped to dynamics of the form (S28) with

$$\begin{aligned} f(\mathbf{u}^E, \theta) &\equiv -\mathbf{u}^E + (1-\alpha)\mathbf{W}^{X\to E}\mathbf{x}, & \text{(S40)} \\ \psi(t) &\equiv \mathbf{W}^{I\to E}\mathbf{Q}\mathbf{c}. & \text{(S41)} \end{aligned}$$

Applied to our parameterization $\mathbf{W}^{X\to E} = \mathbf{M}^E\mathbf{V}^{X\to A}$ and following the least-control framework [87], an update rule for feed-forward parameters $\mathbf{V}^{X\to A}$ can be derived from the surrogate loss $\mathcal{H}(\mathbf{V}^{X\to A}) = ||\mathbf{W}^{I\to E}\mathbf{Q}\mathbf{c}||^2$ as

$$\nabla_{\mathbf{V}^{X\to A}}\mathcal{H}(\mathbf{V}^{X\to A}) = -\left(\frac{\partial f(\mathbf{u}^E, \mathbf{V}^{X\to A})}{\partial \mathbf{V}^{X\to A}}\right)^T \left(\mathbf{W}^{I\to E}\mathbf{Q}\overset{*}{\mathbf{c}}\right) \tag{S42}$$

where $\overset{*}{\mathbf{c}}$ is the control signal at equilibrium that minimizes the loss $\mathcal{L}(\overset{*}{\mathbf{u}})$. From (S37), it is easy to show that the control input at equilibrium can be expressed alternatively as

$$\mathbf{W}^{I\to E}\mathbf{Q}\overset{*}{\mathbf{c}} = \mathbf{W}^{I\to E}\overset{*}{\mathbf{r}}{}^I - \alpha\mathbf{W}^{X\to E}\mathbf{x} \tag{S43}$$

Substituting this expression back into (S42) and evaluating the derivative yields the gradient update rule for the feed-forward parameters $\mathbf{V}^{X \to A}$:

$$\nabla_{v_{kj}^{X\to A}} \mathcal{H}(\mathbf{V}^{X\to A}) = (1-\alpha) \sum_{i}^{N^E} m_{ik}^E \, x_j \left( \alpha \left( \mathbf{W}^{X\to E} \mathbf{x} \right)_i - \left( \mathbf{W}^{I\to E} \mathbf{r}^{*I} \right)_i \right). \tag{S44}$$

To get the BCP update rule defined in the main text (c.f. Eq. (31)), we rewrite the balance deviation in terms of the synaptic currents:

$$\Delta v_{kj} = \frac{\eta_{ff}}{N_A^E} \sum_{i}^{N^E} m_{ik}^E \, x_j \left( \alpha \left( I_i^{exc} - I_i^{inh1} \right)_+ - I_i^{inh2} \right), \tag{S45}$$

where we additionally absorbed the scalar $1-\alpha$ into a small positive learning rate $\eta_{ff}$ and normalized by $N_A^E$ to ensure that the learning rate is independent of the assembly size. This learning rule represents a Hebbian-like learning rule that, for each neuron in the assembly, multiplies presynaptic input by the difference between scaled feed-forward currents and Type 2 inhibitory currents. Note that the formulation in (S45) is strictly speaking non-local, because weights remain tied between all neurons in the assembly. However, the contribution of each neuron remains local and assuming each neuron represents its own assembly, i.e. $N_A^E = 1$, the learning rule reduces to a local Hebbian learning rule.

#### 1.4 Temporally filtered feed-forward currents for online learning

The BCP learning rule derived in Section 1.3 assumes that network dynamics have reached equilibrium for each input before synaptic updates occur. In continuous online learning, however, stimuli evolve over time, and inhibitory currents $I_i^{inh2}(t)$ inevitably lag behind the raw feed-forward drive $I_i^{ff}(t)$ because of finite membrane time constants and recurrent EI dynamics. Here we derive how to low-pass filter $I_i^{ff}(t)$ into a smoothed version $\bar{I}_i^{ff}(t)$ so that it evolves on the same timescale as the network's intrinsic mode—thereby eliminating that lag.

To study the temporal dynamics of E/I assemblies, we consider a linearized system with one excitatory and one inhibitory unit receiving the same feed-forward input:

$$\tau_E \frac{du^E}{dt} = -u^E + I_i^{ff}(t) - w^{I\to E} u^I(t) \tag{S46}$$

$$\tau_I \frac{du^I}{dt} = -u^I + I_i^{ff}(t) + w^{E\to I} u^E(t) \tag{S47}$$

To find the network's intrinsic decay modes, we temporarily set $I_i^{ff}(t) = 0$ and Laplace-transform ($u(t) \mapsto U(s)$):

$$(\tau_E s + 1)\, U^E(s) = -w^{I\to E}\, U^I(s), \tag{S48}$$

$$(\tau_I s + 1)\, U^I(s) = w^{E\to I}\, U^E(s). \tag{S49}$$

Eliminating $U^E$ and $U^I$ yields the characteristic polynomial

$$D(s) = \tau_E \tau_I s^2 + (\tau_E + \tau_I)\, s + \left(1 + w^{I\to E} w^{E\to I}\right) = 0 \tag{S50}$$

whose roots are

$$p_{1,2} = \frac{-(\tau_E + \tau_I) \pm \sqrt{(\tau_E + \tau_I)^2 - 4\tau_E \tau_I (1 + w^{I\to E} w^{E\to I})}}{2\tau_E \tau_I}. \tag{S51}$$

In the regime of moderate coupling, the discriminant remains positive and both $p_1$ and $p_2$ are real and negative. In the limit of weak recurrent gain ($w^{I\to E} w^{E\to I} \ll 1$), these reduce to

$$p_1 \approx -\frac{1}{\tau_E}, \quad p_2 \approx -\frac{1}{\tau_I}. \tag{S52}$$

**Matching the lag via a first-order filter.** Because we care about eliminating the lag—i.e. matching the initial rise of the Type 2 inhibitory feedback—we choose the *faster* of these two modes. Denoting

$$\tau_{\text{adj}} = \min\bigl(\tau_E, \tau_I\bigr), \tag{S53}$$

we approximate the full second-order dynamics by the single leaky integrator

$$\tau_{\text{adj}} \frac{d\bar{I}_i^{\text{ff}}}{dt} = -\bar{I}_i^{\text{ff}}(t) + I_i^{\text{ff}}(t). \tag{S54}$$

This choice ensures that $\bar{I}_i^{\text{ff}}(t)$ rises with the same initial slope as the true inhibitory current $w^{I\to E}u^I(t)$, thereby canceling the leading-edge lag in continuous online learning. Without this temporal alignment, weight updates would be systematically biased, leading to instabilities.

In practice, we have found that setting $\tau_{\text{adj}} = \min(\tau_E, \tau_I)$ yields near-perfect alignment of $\bar{I}_i^{\text{ff}}(t)$ and $I_i^{\text{inh2}}(t)$ across a wide range of membrane time constants. This first-order approximation works well for the input paradigms considered in this paper, where stimuli evolve relatively slowly compared to the network time constants. For different input statistics—such as rapidly varying or oscillatory inputs—alternative approaches may be required, including filters based on the harmonic mean of time constants or full second-order approximations to capture the complete dynamics of the inhibitory response.

#### 2 Supplementary Figures

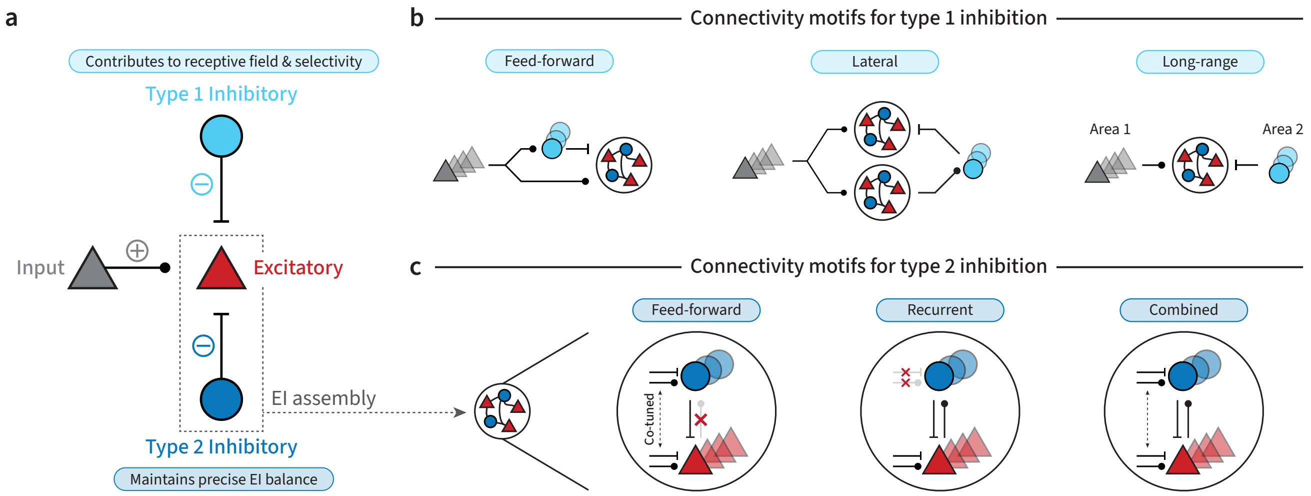

**Figure S1. Functional roles and possible connectivity motifs for Type 1 and Type 2 inhibition.** **(a)** Excitatory neurons in our model receive inhibitory inputs from two types of functionally distinct interneurons: Type 1 (top) and Type 2 (bottom). Type 1 interneurons provide negative contributions to receptive fields and selectivity. Type 2 interneurons form tightly connected EI assemblies with excitatory neurons and maintain a precise EI balance. These functional subtypes are supported by various connectivity motifs. **(b)** Possible connectivity motifs for Type 1 inhibitory interneurons: In feed-forward inhibitory circuits (left), Type 1 interneurons can provide the negative part of a receptive field. Through lateral inhibition (middle), Type 1 interneurons can allow assemblies to inhibit each other. Through long-range inhibition (right), Type 1 interneurons can provide inhibition that is not correlated with excitatory inputs. **(c)** Possible connectivity motifs for Type 2 inhibitory interneurons: Precise balance within an EI assembly can be established with different connectivity mechanisms: If Type 2 interneurons provide feed-forward inhibition (left), they must be co-tuned with the excitatory neurons they maintain a balanced state for. Alternatively, Type 2 interneurons could provide balance through recurrent inhibition (middle), where interneurons provide feedback to the same neurons that excited them. Lastly, feed-forward and recurrent connectivity motifs could be combined to maintain balance (right).

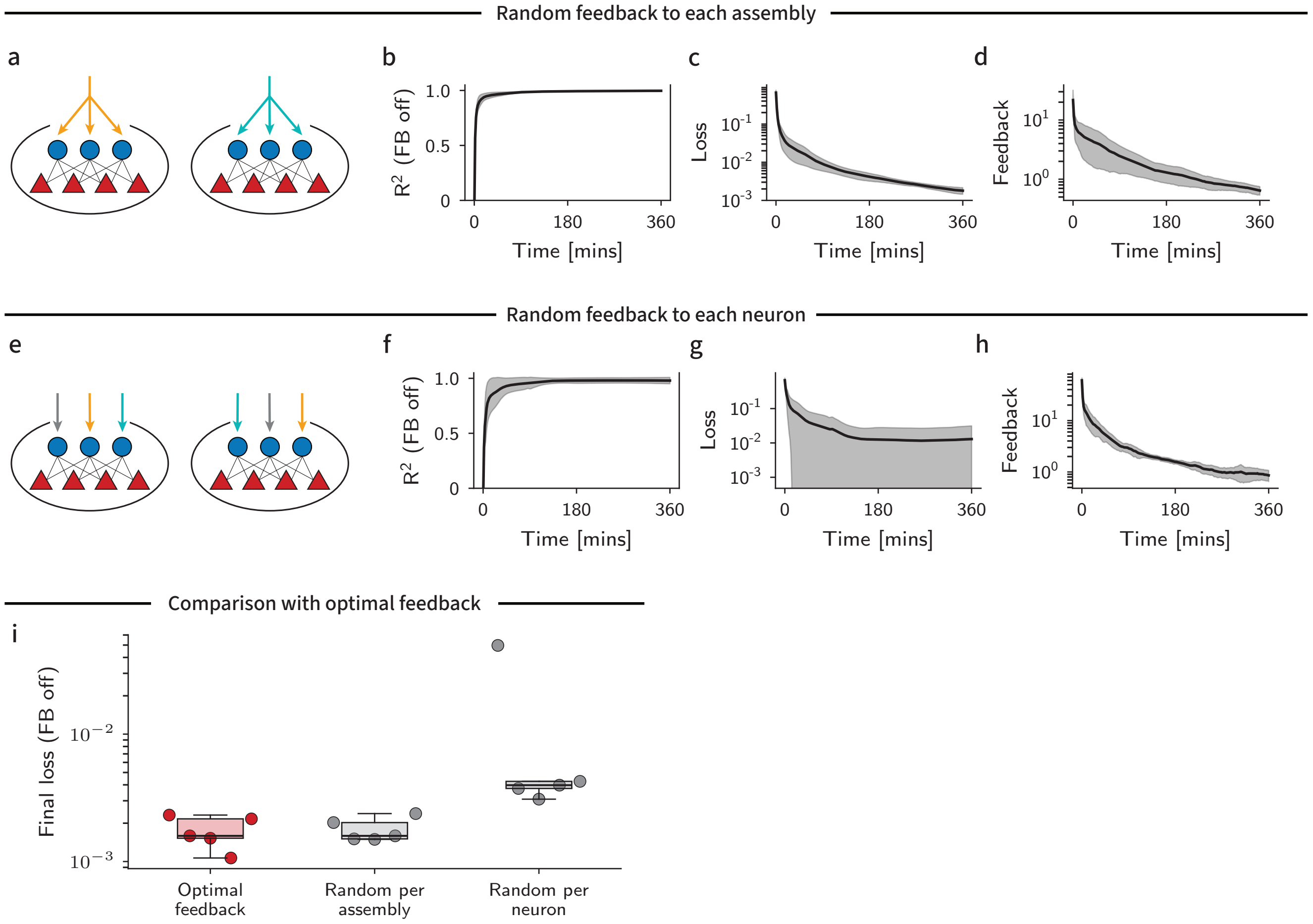

**Figure S2. Online learning in multi-layer networks with random feedback weights.** **(a)** Schematic illustration of assembly-wise random feedback. All Type 2 interneuron in one assembly receive feedback through the same random feedback weights, scaled by each interneurons assembly membership. **(b)** $R^2$ during training of networks with assembly-wise random feedback on the trajectory matching task (c.f. Fig. 4). **(c)** Training loss over training (mean ± SD, n=5 networks). **(d)** Feedback magnitude as a function of time. **(e)** Schematic illustration of neuron-wise random feedback. Top-down feedback to Type 2 interneurons are fully random and do not consider assembly memberships. **(f)-(h)** As panels (c)-(e) for networks with neuron-wise random feedback. **(i)** Final loss in the open-loop (no feedback) setting for networks with optimal feedback weights, assembly-wise random feedback and neuron-wise random feedback.

#### 3 Supplementary Tables

| Parameter | Description | Value |
|---|---|---|
| $N_{\text{E}}$ | Number of excitatory neurons | 16 |
| $N_{\text{I}}$ | Number of inhibitory neurons | 4 |
| $N_{\text{X}}$ | Number of input neurons | 20 |
| $N_{\text{S}}$ | Number of input stimuli | 3 |
| $\tau_{\text{OU}}$ | OU process time constant | 500 ms |
| $\mu_{\text{OU}}$ | OU process mean | 0.0 |
| $\sigma_{\text{OU}}$ | OU process noise | 1.0 |
| $\theta_{\text{OU}}$ | OU process threshold | 0.5 |
| $\tau_{\text{filter}}$ | Stimulus filter time constant | 200 ms |
| $\sigma_X$ | Input neuron tuning width | 0.3 |
| $b_{\text{scale}}$ | Input neuron bias scale | 0.1 |
| $\Delta t$ | Forward euler time step | 1 ms |
| $\tau_{\text{I}}$ | Inhibitory membrane time constant | 20 ms |
| $\tau_{\text{E}}$ | Excitatory membrane time constant | 30 ms |
| $\tau_{\text{out}}$ | Readout time constant | 30 ms |
| $\alpha$ | Precise EI balance parameter | 0.5 |
| $k_p$ | Proportional control gain | 1.0 |
| $k_i$ | Integral control gain | 0.0 |
| $I_+^{\text{ext}}$ | Positive manual feedback strength | 10 |
| $I_-^{\text{ext}}$ | Negative manual feedback strength | $-10$ |
| $\eta_{\text{ff}}$ | Feed-forward learning rate | 0.005 |
| $\tau_{\text{pre}}$ | Pre-synaptic filter time constant | 30 ms |

**Table S1.** Hyperparameters used in simulations of single E/I assemblies (Figs. 2 and 3).

| Parameter | Description | Value |
|---|---|---|
| $N_X$ | Number of input neurons | 20 |
| $N_E$ | Number of excitatory neurons | 240 |
| $N_I$ | Number of inhibitory neurons | 60 |
| $N_A$ | Number of assemblies | 20 |
| $N_Y$ | Number of output neurons | 2 |
| $T_{\text{sin}}$ | Maximum input sinusoid period | 10 s |
| $T_{\text{traj}}$ | Trajectory duration | 5 s |
| $\Delta t$ | Simulation time step | 1 ms |
| $T$ | Total training time | 360 min |
| $N_{\text{iter}}$ | Trajectory iterations | 4320 |
| $\tau_{\text{E}}$ | Excitatory membrane time constant | 30 ms |
| $\tau_{\text{I}}$ | Inhibitory membrane time constant | 20 ms |
| $\tau_{\text{out}}$ | Readout time constant | 30 ms |
| $\alpha$ | Precise EI balance parameter | 0.5 |
| $k_p$ | Proportional control gain | 5.0 |
| $k_i$ | Integral control gain | 0.0 |
| $\eta_{\text{ff}}$ | Feed-forward learning rate | 0.2 |
| $\eta_{\text{out}}$ | Readout learning rate | 0.2 |
| $\tau_{\text{pre}}$ | Pre-synaptic filter time constant | 30 ms |

**Table S2.** Hyperparameters used to train networks on the trajectory matching task (Figs. 4 and 5) and Supplementary Fig. S2.

| Parameter | Description | Value | |
|---|---|---|---|
| | | 1 Hidden Layer | 3 Hidden Layers |
| $N_{\text{X}}$ | Input neurons | 784 | |
| $N_{\text{A}}$ | Assemblies per layer | 128 | |
| $N_{\text{A}}^{\text{E}}$ | Excitatory per assembly | 16 | |
| $N_{\text{A}}^{\text{I}}$ | Inhibitory per assembly | 4 | |
| $N_{\text{Y}}$ | Output neurons | 10 | |
| $B$ | Batch size | 100 | |
| $T_{\text{max}}$ | Max integration time | 2 s | |
| $\tau_{\text{E}}$ | Excitatory membrane time constant | 30 ms | |
| $\tau_{\text{I}}$ | Inhibitory membrane time constant | 10 ms | |
| $\tau_{\text{out}}$ | Readout time constant | 30 ms | |
| $\tau_c$ | Controller time constant | 100 ms | |
| $\eta_{\text{ff}}, \eta_{\text{out}}$ | Learning rate | 0.001 | |
| $\alpha$ | Precise EI balance parameter | 0.5 | |
| $k_p$ | Proportional control gain | 0.2 | 0.05 |
| $k_i$ | Integral control gain | 0.4 | 0.1 |

**Table S3.** Hyperparameters used in training networks on the Fashion-MNIST dataset (Fig. 6). Different control parameters were used for 1-layer and 3-layer networks.

| Parameter | Description | Value |
|---|---|---|
| $N_X$ | Number of input neurons | 20 |
| $N_E$ | Number of excitatory neurons | 160 |
| $N_I$ | Number of inhibitory neurons | 40 |
| $N_A$ | Number of assemblies | 40 |
| $N_Y$ | Number of output neurons | 1 |
| $\Delta t$ | Simulation time step | 1 ms |
| $\tau_{\text{OU}}$ | Noise OU process time constant | 200 ms |
| $\mu_{\text{OU}}$ | Noise OU process mean | 1.0 |
| $\sigma_{\text{OU}}$ | Noise OU process variance | 0.05 |
| $\tau_{\text{rise}}$ | Rise time of the input stimulus | 40 ms |
| $\tau_{\text{decay}}$ | Decay time of the input stimulus | 200 ms |
| $\tau_{\text{E}}$ | Excitatory membrane time constant | 20 ms |
| $\tau_{\text{I}}$ | Inhibitory membrane time constant | 10 ms |
| $\tau_{\text{out}}$ | Readout time constant | 20 ms |
| $\theta$ | Freezing readout shift | -0.6 |
| $a$ | Freezing readout steepness | 4 |
| $\alpha$ | Precise EI balance parameter | 0.5 |
| $k_p$ | Proportional control gain | 1.0 |
| $k_i$ | Integral control gain | 0.0 |
| $\eta_{\text{ff}}$ | Feed-forward learning rate | 10.0 |
| $\eta_{\text{out}}$ | Readout learning rate | 0.0 |
| $\tau_{\text{pre}}$ | Pre-synaptic filter time constant | 100 ms |

**Table S4.** Hyperparameters used in training networks on a task mimicking fear conditioning involving the BLA (Fig. 7).

| Parameter | Description | Value |
|---|---|---|
| $N_X$ | Number of input neurons | 20 |
| $N_E$ | Number of excitatory neurons | 320 |
| $N_I$ | Number of inhibitory neurons | 80 |
| $N_A$ | Number of assemblies | 80 |
| $N_Y$ | Number of output neurons | 1 |
| $\Delta t$ | Simulation time step | 1 ms |
| $\tau_{\text{OU}}$ | Noise OU process time constant | 500 ms |
| $\mu_{\text{OU}}$ | Noise OU process mean | 0.0 |
| $\sigma_{\text{OU}}$ | Noise OU process variance | 0.05 |
| $N_{\text{sines}}$ | Number of sine waves for input generation | 10 |
| $\tau_{\text{rise}}$ | Rise time of the cue-evoked input | 30 ms |
| $\tau_{\text{decay}}$ | Decay time of the cue-evoked input | 500 ms |
| $\tau_{\text{E}}$ | Excitatory membrane time constant | 20 ms |
| $\tau_{\text{I}}$ | Inhibitory membrane time constant | 10 ms |
| $\tau_{\text{out}}$ | Readout time constant | 50 ms |
| $\alpha$ | Precise EI balance parameter | 0.5 |
| $k_p$ | Proportional control gain | 10.0 |
| $k_i$ | Integral control gain | 2.0 |
| $\tau_c$ | Controller time constant | 100 ms |
| $\eta_{\text{ff}}$ | Feed-forward learning rate | 0.02 |
| $\eta_{\text{out}}$ | Readout learning rate | 0.01 |
| $\tau_{\text{pre}}$ | Pre-synaptic filter time constant | 50 ms |

**Table S5.** Hyperparameters used in training networks on a task mimicking motor learning involving M1 (Fig. 8).
